# DROSHA, DICER and Damage-Induced long ncRNA control BMI1-dependent transcriptional repression at DNA double-strand break

**DOI:** 10.1101/2024.08.07.606960

**Authors:** Francesca Esposito, Ilaria Capozzo, Stefania Modafferi, Ubaldo Gioia, Letizia Manfredi, Fabio Iannelli, Alessio Colantoni, Adelaide Riccardi, Alessia di Lillo, Sara Tavella, Matteo Cabrini, Fabrizio d’Adda di Fagagna, Sofia Francia

## Abstract

Genome integrity is safeguarded by the DNA damage response (DDR). Controlled transcriptional dampening of genes surrounding DNA double-strand breaks (DSBs) has been shown to facilitate DNA repair. This phenomenon, defined as DSB-induced silencing in *cis* (DISC), involves the DDR apical kinase ATM and the Polycomb Repressive Complex 1 (PRC1). Conversely, DSBs have also been reported to induce *de novo* transcription of damaged-induced long non-coding RNAs (dilncRNAs) in a MRE11-RAD50-NBS1 (MNR) complex-dependent manner. MRN also controls the recruitment to DSB of the ribonuclease DROSHA, which together with DICER, stimulates DDR signaling and DNA repair. Here, we reconcile these apparently contrasting observations by showing that dilncRNA, together with DROSHA and DICER, but not GW182-like proteins required for miRNA-mediated gene silencing, controls DISC. Indeed, similarly to ATM, MRN inhibition abolishes DISC while pharmacological enhancement of DICER ribonuclease activity by Enoxacin improves DISC. Importantly, Enoxacin administration restores DISC upon ATM inhibition, demonstrating that DICER promotes DISC independently from ATM. Differently, Enoxacin does not restore DISC upon MRN inhibition, suggesting that DICER acts downstream to dilncRNA biogenesis and DROSHA recruitment. Mechanistically, we show that DROSHA and DICER control the recruitment of the PRC1 component BMI1 at DSBs and the consequent H2A-K119 ubiquitination. Upon DSBs formation, BMI1 and DROSHA interact in an RNA-dependent manner. Indeed, BMI1 associates to dilncRNA and do so in a DROSHA- and DICER-dependent manner. Importantly, inhibition of dilncRNA function by antisense oligonucleotides or Cas13-mediated targeting is sufficient to reduce BMI1 recruitment and DISC at individual loci. We propose that dilncRNAs together with DROSHA and DICER control DISC at genomic DSB by supporting PRC1 recruitment and chromatin ubiquitination.

## Introduction

At DNA damage sites, generation of RNA-interference (RNAi) machinery-dependent non-coding RNAs has been observed in several organisms and different genomic contexts ^1–5^. In mammalian cells, we and others have observed that a double strand break (DSB) induces *de novo* transcription of non-coding RNAs ^6–10^. We proposed that DSBs recruit the DNA-damage’s sensor complex MRE11-RAD50-NBS1 (MRN) that, by unwinding DNA ends^11^, allows the loading and the activity of RNA polymerase II (RNAPII) together with its Pre-Initiation Complex (PIC) ^12^. This leads to the *de novo* transcription of damage-induced long non-coding RNAs (dilncRNAs) ^6,7,11,12^. Concomitantly, also DROSHA ribonuclease is recruited to DNA ends ^13,14^, suggesting that transcripts can be locally processed by DROSHA at DSB. DROSHA recruitment occurs in a MRN-dependent —but ATM-independent— manner and promotes 53BP1 recruitment and DNA repair by Non-Homologous End-Joining (NHEJ)^13,14^. Others, have shown that upon DNA damage, phosphorylated DICER migrates into the nucleus where it associate with DSB ^15,16^. Thus, DROSHA and DICER subsequent RNA processing at DSB can generate DNA-damage RNAs (DDRNAs) that together with dilncRNAs promote the activation of the DNA damage response (DDR) pathway ^17–21^. DDRNAs are mostly produced at repetitive loci and indeed their biogenesis occurs also at dysfunctional telomeres ^22,23^. DROSHA has also been proposed to mediate the formation of DNA:RNA hybrids at DSBs occurring in transcribed endogenous loci, supporting the notion that the RNAi-machinery is present and active at DSBs ^14^.

Concurrently, we and others have also shown that transcription of coding genes in proximity of a DSB is actively repressed through different mechanisms ^24–32^. Specifically, kinase activity of ATM, an apical DDR kinase, has been causally involved in repression of transcription at DSBs, also known as DSB-induced silencing *in cis* (DISC) ^24,25,29,33^. ATM phosphorylation of Brahma-related gene-1 (BRG1) and hbrm-associated factor 180 (BAF180), a Polybromo-associated BAF (PBAF) remodeling complex subunit, has been reported to induce DISC ^26^. Polycomb group complexes 1 and 2 (PRC1 and PRC2) have been shown to be required for DISC ^26^. In addition, ATM-dependent phosphorylation of ENL, a component of the elongating transcriptional machinery, stimulates its association with the PRC1 subunit, BMI1. This interaction promotes the accumulation of BMI1 at actively transcribed genes flanking a DSB and the consequent PRC1-mediated mono-ubiquitylation of histone H2A on lysine 119 (ubH2A-K119), thus promoting chromatin compaction and DISC in proximity to DSBs ^27^. BMI1-mediated ubH2A-K119 has also been reported to promote DNA repair ^34^.

Therefore, DSBs induce both *de novo* transcription of non-coding RNAs, which can be processed by the components of the RNAi machinery DROSHA and DICER, as well as repression of active, canonical genes positioned nearby. How these two apparently contradictory transcriptional events influence each other remains unknown.

Here, we demonstrate that DROSHA, DICER and dilncRNAs control DSB-induced transcriptional silencing of surrounding genes. Mechanistically, we show in different cellular systems and genomic context that DROSHA and DICER, together with dilncRNAs stimulate BMI1 recruitment at DSB and UbH2A-K119 deposition, thus controlling DISC. Notably, precise dilncRNAs targeting via both antisense oligonucleotides (ASOs) or Cas13 nuclease impairs DISC at individual loci, supporting a role for DSB-induced RNAs in this mechanism.

## Results

### DICER is required for DISC of a reporter gene proximal to a cluster of DSBs

In *Drosophila melanogaster* and *Neurospora crassa* cells, DNA damage-induced and RNAi-dependent small ncRNAs have been proposed to act as endogenous siRNAs, silencing genes encoded by the region flanking DNA lesions ^1,35^. We thus hypothesized that DROSHA and DICER’s RNA products may act in a similar fashion, repressing transcription of active genes flanking DSBs in mammalian cells. We therefore investigated the potential contribution of DROSHA, DICER in DISC. To test this hypothesis, we initially took advantage of the engineered U2OS-2-6-3 cellular system previously created to study DISC and widely used by several other research groups ^24,26,27,36^. In this system, a cluster of DSBs is generated upstream of an inducible reporter gene and flanked by hundreds of copies of Lac operator (LacO). Thus, the expression of the FokI endonuclease fused to a Lac inhibitor (LacI) and the fluorescent mCherry tag allows for both the generation of multiple DSBs at the LacO repeats and the visualization of the genomic locus in the nucleus (Figure 1A) ^24^. Nuclease-deficient D450A FokI mutant (FokI D450A) is used as negative control for DSB generation, while still marking the target locus under undamaged conditions. Downstream of the cleavable repeats, cells bear the gene encoding for a CFP-tagged peroxisomal targeting peptide (CFP-SKL), containing 24 stem-loops that are bound by the yellow fluorescent protein (YFP)-tagged MS2 protein, expressed upon doxycycline administration (Figure 1A). Thus, YFP-MS2 binding to nascent CFP-SKL transcripts allows for the visualization of RNA generation ^24^. In the same system, a precise quantification of nascent transcripts can be achieved by RT-qPCR analysis at the CFP-SKL gene. Overall, this is a well-established cellular system to monitor DISC.

**Figure 1.**
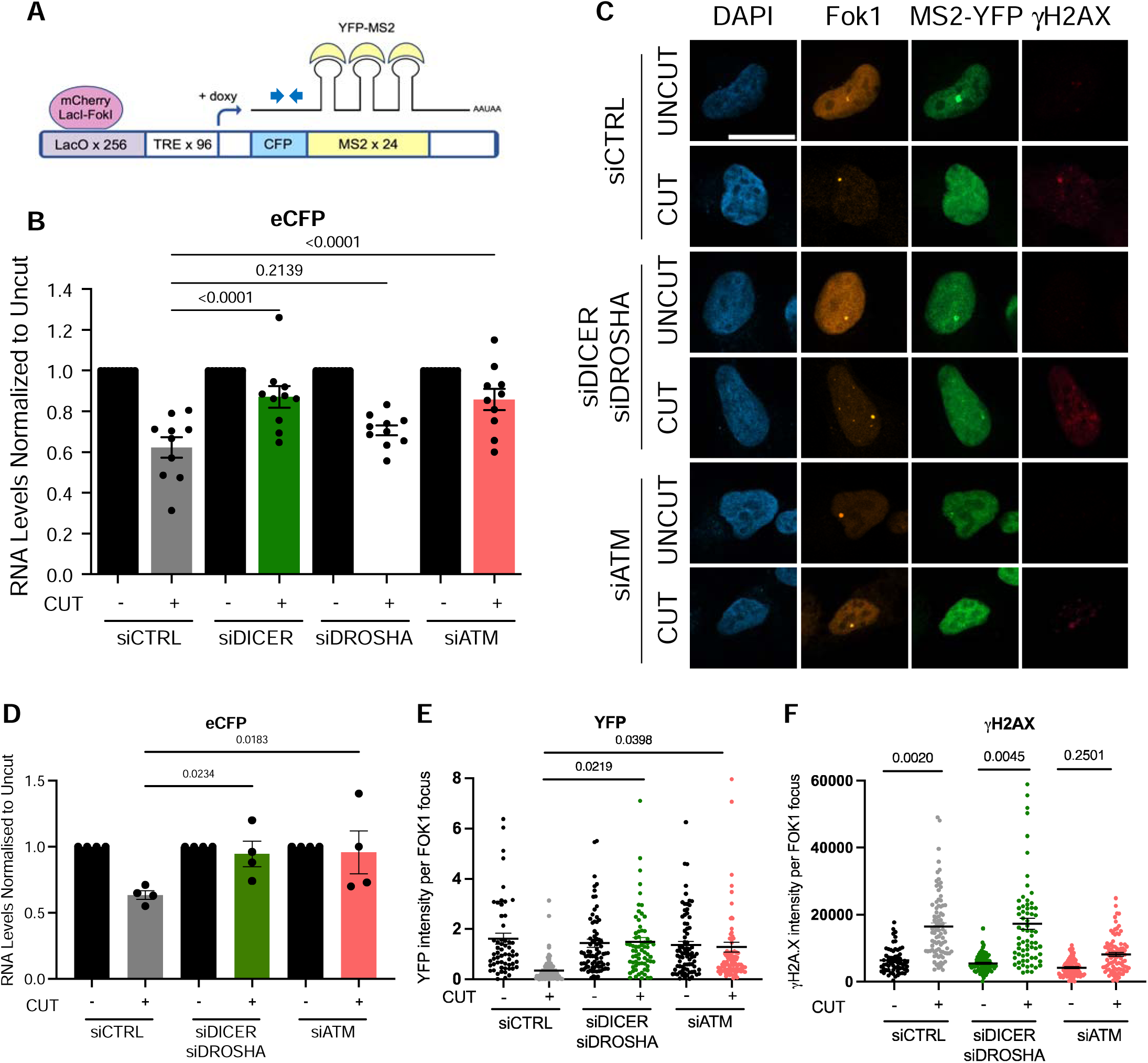
DICER and DROSHA are required for DISC in U2OS 2-6-3 cells. **A.** Schematic representation of the reporter locus in U2OS 2-6-3 cells. **B.** RT-qPCR analysis of CFP mRNA levels in uncut (mutant D450A FOK1 transfected) and cut cells (WT FOK1 transfected), treated with non-targeting siRNAs (siCTRL) or siRNAs against DICER (siDICER), DROSHA (siDROSHA) and ATM (siATM) transcripts. Error bars represent SEM from eleven independent experiments. Values are relative to uncut cells for each knockdown condition. **C.** Representative images of YFP-MS2 and γH2AX signals at the Cherry-Lac locus in uncut and cut U2OS 2-6-3 cells transfected with siCTRL, siDICER and siDROSHA, or siATM. Scale bar: 10 µm. **D.** RT-qPCR analysis of CFP mRNA levels in cells treated as in C. Error bars represent SEM from four independent experiments. Data are relative to uncut cells for each knockdown condition. **E.** Quantification of YFP-MS2 fluorescence intensity in cells treated as in C. Error bars represent SEM of more than 50 cells from three independent experiments. **F.** Quantification of γH2AX fluorescence intensity in cells treated as in C. Values are relative to uncut cells for each knockdown condition. Error bars represent SEM of more than 50 cells from three independent experiments. Statistical test used in B, D, E and F was One-Way ANOVA.

In this system expression of Wild-Type (WT) Fok1, but not of mutant D450A Fok1, reduces CFP-SKL transcript levels by 40-50% as detected by RT-qPCR (Figure 1B), and ATM knock-down prevents CFP-SKL silencing (Figure 1B), as previously reported ^24^. With the same approach, we tested the impact of DROSHA or DICER knockdown on DISC. We observed that DICER knockdown dramatically abolishes DISC (Figure 1B), while DROSHA knockdown alone showed only modest impact on DISC despite similar knockdown efficiency (Supplementary Figure 1A). This suggests a stronger role for DICER compared to DROSHA in DISC when multiple DSBs are generated in a cluster at repetitive sequences upstream of a gene.

We and others observed that the Translin/Trax complex can compensate the loss of DICER’s activity, but it cannot act if also DROSHA is concomitantly inactivated ^6,37^. Therefore, we extended our observations by combining DROSHA and DICER inactivation (co-KD), prior to DISC evaluation. Indeed, the results obtained by both RT-qPCR analysis of the CFP-SKL mRNA levels and by the quantification of the intensity of the YFP-MS2 fluorescent signal, clearly demonstrates that the concomitant depletion of DROSHA and DICER, dramatically abolishes DISC to the same extent of ATM inactivation (Figure 1C-E). Importantly, the variations observed in DISC were not due to different levels of DNA damage, since immunofluorescence (IF) for γH2AX demonstrated a similar amount of damage in all cut samples, as previously observed in other cellular systems ^6,17,20^ (Figure 1F). Importantly, co-KD of DROSHA and DICER strongly reduces histone H2A mono-ubiquititation of lysin 119 (ubH2A-K119), the chromatin modification associated to DISC (Figure S1C,D). Consistently with previous observations in this system, ATM inactivation led to a decrease in both γH2AX and ubH2A-K119 ^24,26,27,36^ (Figure 1F and S1C,D). Based on these results, we preferentially used concomitant inactivation of DROSHA and DICER as a more efficient approach to study the role of these two RNases in DISC.

Altogether, this set of data indicates that DICER is required for transcriptional silencing of genes next to heavily damaged chromatin and does so to the same extent as ATM when is also supported by DROSHA.

### DROSHA and DICER control DISC of endogenous genes caused by individual DSBs

To extend and strengthen our conclusions at endogenous genomic loci, we took advantage of the DIvA cellular system ^38^. In these cells, the restriction enzyme AsiSI is expressed under the control of a modified version of the Estrogen-Receptor binding domain, thus its nuclear localization and DNA cleavage activity can be induced by the addition of 4-Hydroxytamoxifen to the medium ^38,39^. By cap analysis of gene expression (CAGE) in this cellular system, we previously reported that DISC also takes place at endogenous genes and that ATM inhibition prevents it ^29^. Here, we performed RNA sequencing in cells co-KD for DROSHA and DICER, both in cut and uncut conditions, for five independent experiments. Non-targeting siRNA-transfected cells or ATM KD cells were used as negative and positive control for DISC, respectively. In our computational analyses we considered the expression level of genes previously reported to be reproducibly cut in AsiSI sites within 2Kb from gene bodies ^29^. Among these, we focused on the 98 genes that were consistently downregulated upon cut in control samples (siCTRL). Relative differential expression analyses of the 98 genes, revealed that DROSHA and DICER co-KD reduces DISC similarly to ATM depletion, as indicated by reduced fold-change in genes’ expression level between cut and uncut conditions (Figure 2A and S2A). Therefore, DROSHA and DICER control DISC also at endogenous genes upon induction of individual DSBs at different, non-repetitive, genomic loci.

**Figure 2.**
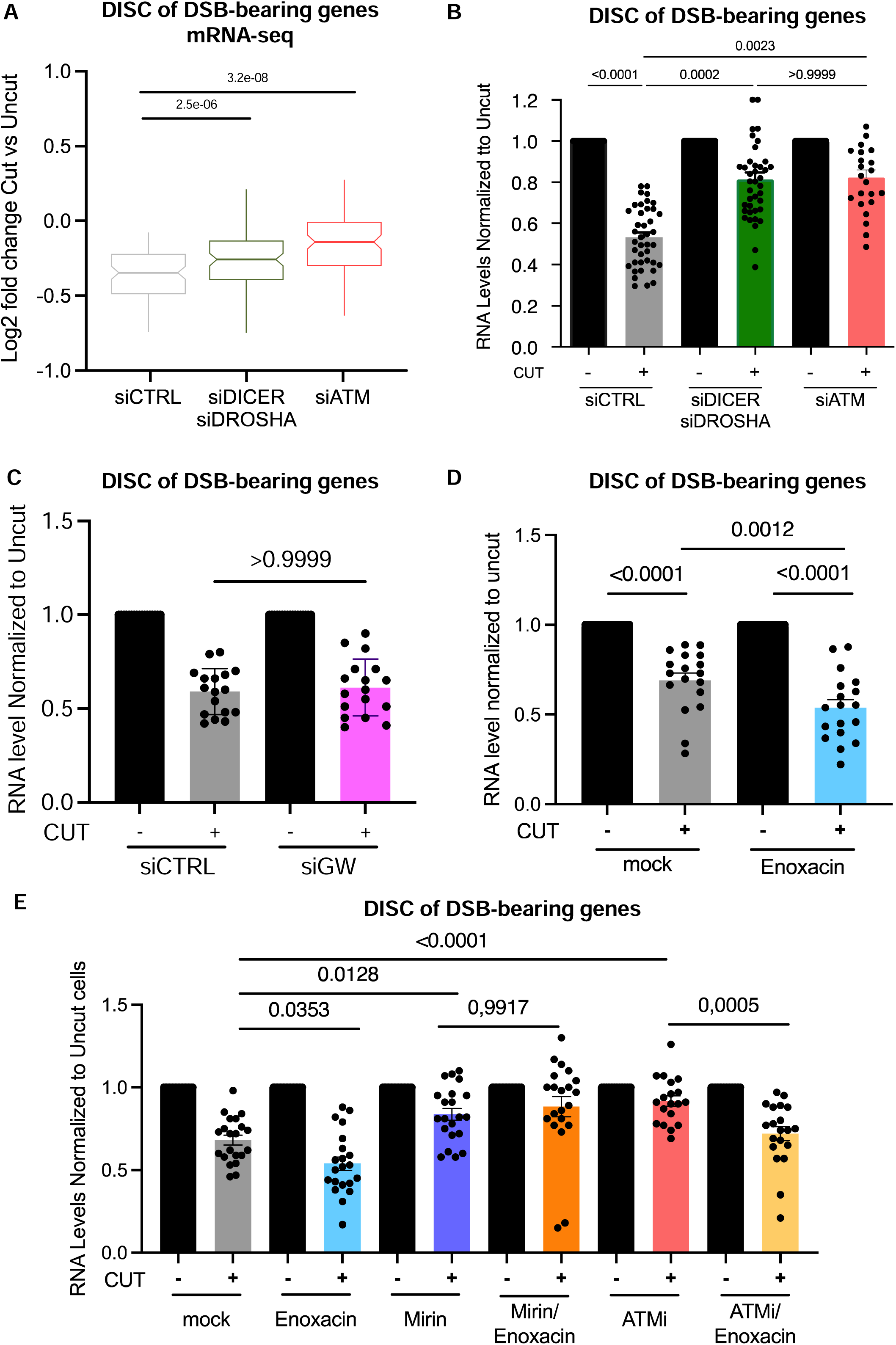
DISC is controlled by DROSHA and DICER in DIvA cells. **A.** Box plot representing the fold change of transcript levels of genes in proximity to AsiSI sites compared to uncut condition, in cells transfected with non-targeting siRNAs (siCTRL), with siRNAs targeting DROSHA and DICER (siDICER, siDROSHA) or ATM (siATM); 98 genes bearing AsiSI sites, within or adjacent (±2 Kb) to gene bodies, that underwent downregulation upon cut in all the three independent biological replicates, are shown. Statistical analysis was performed using the paired Wilcoxon test. **B.** The expression of a subset of 6 break-bearing genes was validated by RT-qPCR in cells treated as in A. Data are relative to uncut cells for each knockdown condition. **C.** The expression of the 6 break-bearing genes was studied by RT-qPCR in DIvA cells treated with either siCTRL or siRNAs against GW-like complex subunits TNRCA, TNRCB or TNRCC (siGW A, B, C). Values are relative to the siCTRL uncut condition. **D**. The mRNA levels of the selected genes were analysed by RT-qPCR in cells treated or not with enoxacin. Data are relative to the untreated and uncut condition. **E.** The expression of each selected gene was analysed by RT-qPCR in samples treated or not with enoxacin, KU-60019 (ATMi) and Mirin. Values are relative to untreated and uncut samples for each condition. Statistical analyses in panels B-E were performed using One-Way ANOVA. Error bars represent SEM of the expression of the 6 selected genes from three independent experiments.

Top-ranked cut sites in the DIvA cellular system were selected over the years crossing γH2AX ChIP-seq signals and Breaks Labeling In Situ and Sequencing (BLISS) data set in our previous studies ^13,29^. To validate the RNA-sequencing data at individual genomic sites, we selected 6 genes bearing cut AsiSI sites at different positions with respect to the adjacent gene bodies, [*i.e.*: TRIM37, RBMXL1 and HUNK genes are cut in promoters or at the Transcriptional Start Site (TSS); GNE and MIS12 have DSB within the gene body; KLF7 gene is cut in its 3’UTR] and tested the impact of DROSHA and DICER inactivation on their DISC. By RT-qPCR, we observed that DSB induction dampened the expression of all these genes and that DROSHA and DICER co-KD sustained transcription despite proximal DNA damage (Figure 2B and S2A). Additionally, individual KD of DICER or DROSHA was sufficient to reduce DISC (Figure S2B and S2C). Of note, DROSHA and DICER co-KD did not impact on the expression levels of the same genes in the uncut condition (Figure S2D), suggesting that miRNAs are not implicated in the DISC observed. To more thoroughly demonstrate that, we simultaneously knocked down the three GW182-like proteins (TNRCA, -B or -C, Figure S2E) which play a central role in miRNA-mediated gene silencing ^40^, but not in DDR ^17^. Importantly, we observed unaltered DISC in cells co-KD for TNRCA, B and C (Figure 2C). To further exclude a role for miRNA loss in our settings, we expressed the TNRCA peptide fragment T6B, previously reported to interfere with Argonaute protein assembly in the RISC complex ^41^, or a mutant and inactive form as control. In these same DSB-bearing genes we observed by RT-qPCR unaltered gene expression upon T6B peptide expression, (Figure S2F), altogether demonstrating that DROSHA end DICER activity in miRNA generation is unlikely to play a significant contribution to DISC.

In this DIvA cellular system, a subset of AsiSI sites have been shown to be repaired exclusively by non-homologous end-joining (NHEJ-prone), while others are preferentially repaired by homologous recombination (HR-prone sites) ^39^. We have previously reported that upon DNA damage induction, DROSHA is recruited to both NHEJ- and HR-prone sites ^13^. To test if DROSHA and DICER control DISC at either NHEJ-or HR-prone DSBs or both, we compared DISC of genes in the two subgroups and the impact that DROSHA and DICER co-KD had on it. We observed that all genes were downregulated to the same extent upon cut, and that DROSHA/DICER co-KD abolished DISC in both NHEJ-prone and HR-prone DSBs (figure S2G).

The antibiotic Enoxacin has been shown to stimulate the endoribonuclease activity of the DICER complex by facilitating the interaction between the HIV TAR RNA binding protein (TRBP) and its RNA substrates ^42^. Recent work from our group demonstrated that Enoxacin administration at doses that do not alter miRNA levels, boosts DDR signaling and DNA damage repair via NHEJ by increasing DICER-dependent DDRNA biogenesis ^43^.

Therefore, we tested if Enoxacin could boost DISC. To do so, DIvA cells were treated with Enoxacin before induction of damage, and DISC was measured by RT-qPCR at the previously characterized DSB-bearing genes. Intriguingly, Enoxacin treatment significantly enhanced DISC at DSB-containing genes (Figure 2D), while leaving gene expression unaltered in uncut condition (Figure S2H), suggesting that local DDRNA production upon damage promotes DISC at DSB-bearing genes. Enoxacin ability to boost DDR ^43^ was confirmed in these settings by the observed higher numbers of 53BP1 foci in treated DIvA cells (Figure S2I), without altering cell-cycle phase distribution (Figure S2J). These results indicate that Enoxacin, by stimulating DDRNA production and DDR signaling, strengthens DISC of DSB-bearing genes, further confirming the functional involvement of DDRNAs in this mechanism.

The MRN complex controls both ATM activation ^44^ and the recruitment of RNAPII ^6,11,12^ and DROSHA ^13^ at DSBs. Therefore, to further validate the role of both ATM and DROSHA in DISC, we tested if treatment with the MRN-complex inhibitor, mirin, could abrogate DISC. As expected, mirin administration completely restored transcription of break-bearing genes (Figure S2K). Since DICER and DROSHA sustain ATM activation ^6,17^, we wondered whether they promote DISC via the ATM’s kinase activity or throughout a distinct parallel mechanism, too. To do so, we tested if Enoxacin could restore DISC upon pharmacological inhibition of ATM’s kinase activity. As previously shown^29^, ATM inhibition abolished DISC in the DIvA cellular system (Figure 2E). Importantly, we observed that Enoxacin treatment significantly repressed transcription of damaged genes in cells in which ATM’s activity was inhibited (Figure 2E), demonstrating that DICER acts in DISC also independently from ATM. Conversely, in these same settings, MRN inhibition by mirin abolished Enoxacin ability to stimulate DISC (Figure 2E). This result suggests that Enoxacin acts only in an MRN-dependent manner in DISC, consistently with the fact that MRN activity -but not ATM one-is required for dilncRNA synthesis and DROSHA recruitment to DNA-damage sites ^6,11–13^. On the other hand, there is no evidence connecting MRN activity to microRNA processing, further supporting a model where Enoxacin acts in DISC independently from microRNA.

DNA damage can occur in different forms. To extend our observations beyond DSB induced by restriction enzymes, we generated DNA damage in a more complex and site-independent fashion by laser micro-irradiation, and studied DISC by pulsed EU incorporation as previously performed in ^45^ and ^33^. After DROSHA and DICER co-KD (Figure S2K), we laser micro-irradiated small areas of the nuclei and exposed cells to 1-hour pulse of EU to label newly synthesized RNA. We measured DISC by detecting EU signals by click chemistry in areas of DNA damage determined by immunofluorescence (IF) against γH2AX. Also in this additional setting, we observed transcriptional silencing at chromatin damaged by laser micro-irradiation (Figure S2L-M). DROSHA and DICER co-KD caused a higher EU signal in γH2AX-positive areas compared to control siRNA (siCTRL)-treated cells (Figure S2L-M). Therefore, DROSHA and DICER are key for DISC also at DNA lesions generated by laser micro-irradiation.

Altogether, these results indicate that DROSHA- and DICER-mediated DDR signaling has a strong impact on transcriptional control of genes adjacent to DSB generated by different means.

### DROSHA and DICER control BMI1 recruitment and chromatin ubiquitination at DSB

PRC1-mediated mono-ubiquitylation of histone H2A on lysine 119 (ubH2A-K119) is known to induce chromatin compaction and transcriptional repression in different contexts ^46^, and contribute to DNA repair ^34^. Previous work in the U2OS 2-6-3 cellular reporter system indicates that upon generation of multiple DSBs, DISC is enforced by ATM via the phosphorylation of the transcription elongation factor ENL, an event that stimulates the recruitment of the PRC1 component and E3-ubiquitin ligase BMI1 to the reporter gene ^27^. Intriguingly, BMI1 has also been shown to physically interact with the DDR sensor NBS1 ^34,47^, a component of the MRN complex. In addition, BMI1 is recruited to laser tracks in an ATM, γH2AX and RNF8 dependent manner, and loss of BMI1 sensitizes cells to DNA damaging agents ^34^. Finally, BMI1 has been recently involved in DNA-end resection and repair by HR ^48^. These observations suggest a model in which PRC1 is actively recruited to the DSB where it ubiquitinates chromatin leading to DISC, thus favoring DNA repair. However, the recruitment of BMI1 to individual DSBs in the DIvA cells is controversial ^49^. To test the role of BMI1 in DISC of DSB-bearing genes, we knocked-down BMI1 in DIvA cells and measured the level of transcription of the set of genes previously characterized. BMI1 KD inhibited DISC to an extent similar to the one observed upon ATM KD (Figure 3A and S3B) demonstrating that BMI1 controls DISC also in DIvA cells. To test the role of DROSHA/DICER in H2A-K119 ubiquitination at DSBs, we stained cells KD for BMI1, DROSHA/DICER or ATM, with antibodies against ubH2A-K119 and γH2AX. Indeed, BMI1 and DICER/DROSHA co-KD significantly reduced the formation of detectable ubH2A-K119 foci, while leaving γH2AX foci and H2AX protein levels unaltered (Figure 3B-D, S3A) – as expected, ATM-depleted cells showed a significant reduction in γH2AX signal too (Figure 3B and 3C). Of note, neither DICER/DROSHA nor BMI1 KD significantly altered ubH2A-K119 signal in undamaged cells (Figure 3D). These results support the notion that BMI1 controls chromatin ubiquitination at individual DSBs, and it is required for DISC or DSB-bearing genes in DIvA cells.

**Figure 3:**
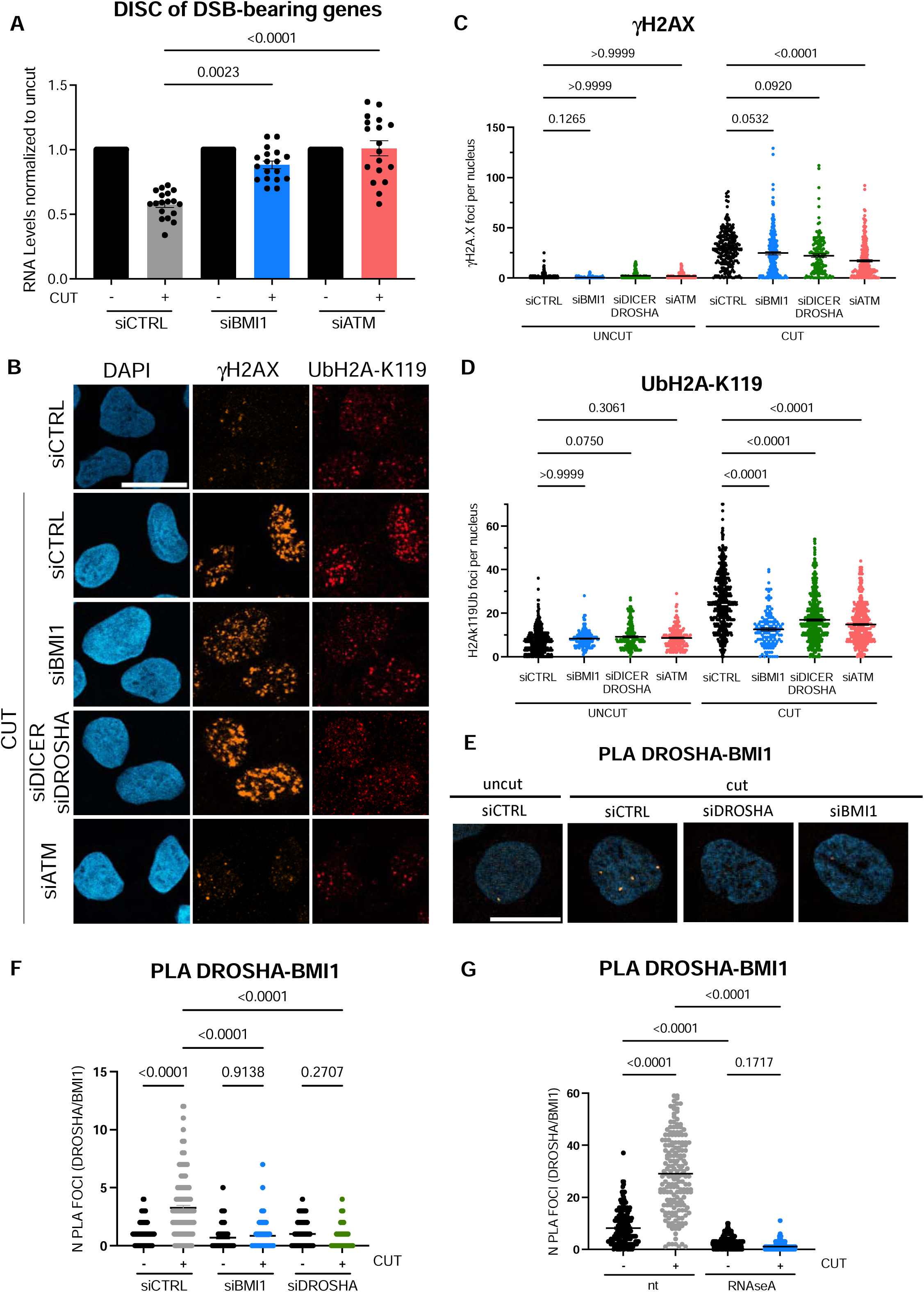
BMI1, DICER and DROSHA control ubH2A-K119 deposition in DIvA cells; RNA controls the proximity between BMI and DROSHA induced in damaged cells. **A.** The mRNA levels of 6 different break-bearing genes, from samples transfected with non-targeting siRNAs (siCTRL) or siRNAs against BMI1 (siBMI1) or ATM (siATM) transcripts, were analysed by RT-qPCR. Data are relative to uncut cells for each knockdown condition. Error bars represent SEM of the expression of the selected 6 genes from three independent experiments. **B.** Images of cells transfected with siCTRL, siBMI1, siDICER and siDROSHA or siATM and stained for γH2AX and ubH2A-K119. DNA was counterstained with DAPI. Scale bar: 20 µm. **C.** Quantification of γH2AX foci in cells treated as in B. Error bars represent SEM of more than 150 cells from three independent experiments. **D.** Quantification of ubH2A-K119 foci in cells treated as in B. Error bars represent SEM of more than 150 cells from each of the three independent experiments. **E.** Representative images of BMI1-DROSHA PLA signals in uncut and cut DIvA cells treated with siCTRL, siDROSHA or siBMI1. Scale bar: 20 µm. **F.** Quantification of BMI1-DROSHA PLA signal in cells treated as in E. Error bars represent SEM of more than 150 cells from each of the three independent experiments. **G.** Quantification of BMI1-DROSHA PLA signal in uncut and cut DIvA cells non treated (nt) or treated with RNAseA after PFA fixation. Statistical analyses in panels A, C, D, F and G were performed by One-Way ANOVA.

To further clarify the relative contribution of BMI1-dependent ubH2A-K119 deposition on ATM activation, we tested whether BMI1-KD might negatively impact on ATM activation and DDR foci formation at DSB. To do so, we stained for γH2AX, 53BP1 and pATM in both cut and uncut DIvA cells upon KD of BMI1 (Figure S3B-H): we confirmed that BMI1 is not required for γH2AX and 53BP1 foci formation (Figure S3C-F) and, rather, its loss seemed to stimulate ATM activation (Figure S3G,H) as previously proposed ^50^.

To seek for a mechanism of BMI1 recruitment we tested if DROSHA and BMI1 could display vicinity upon DNA damage in a Proximity Ligation Assay (PLA). PLA signal obtained using primary antibodies against DROSHA and BMI1 was induced upon DNA damage (Figure 3E and 3F). The signal is lost in DROSHA and BMI1 KD cells, demonstrating its specificity. To test weather this interaction was mediated by RNA, we treated cells with RNase-A prior to PLA assay. Importantly, PLA signals were completely abolished upon RNase-A treatment, suggesting that BMI1-DROSHA proximity is mediated by RNA (Figure 3G).

Thus, we hypothesized that DROSHA, DICER and dilncRNA might control chromatin ubiquitination at sites of DNA damage and DISC by promoting BMI1 recruitment. To test this hypothesis, we performed a chromatin immunoprecipitation followed by qPCR (ChIP-qPCR) analysis for BMI1, ubH2A-K119 and γH2AX in control or DROSHA/DICER-co-KD conditions (Figure 4A,B and S4A,B). Three different DSBs (DS1, DS2 and DS3) were analyzed, while an unrelated region far from AsiSI sites was used as a negative control. At all DSBs tested, we observed a strong recruitment of BMI1, which was lost upon DROSHA and DICER co-KD (Figure 4A and S4A). Moreover, the lack of BMI1 recruitment was comparable to the one observed upon ATM KD, further strengthening the key role of DROSHA and DICER factors in DISC. ChIP-qPCR analyses for ubH2A-K119 and γH2AX at the same three AsiSI sites (DS1, DS2 and DS3) revealed a strong accumulation of ubH2A-K119 signal upon DNA damage, which was partly ATM-dependent (Figure 4B). Similarly, DROSHA and DICER co-KD significantly reduced the accumulation of ubH2A-K119 signal on damaged chromatin as detected by ChIP-qPCR (Figure 4B), while leaving γH2AX unaffected (Figure S4B).

**Figure 4:**
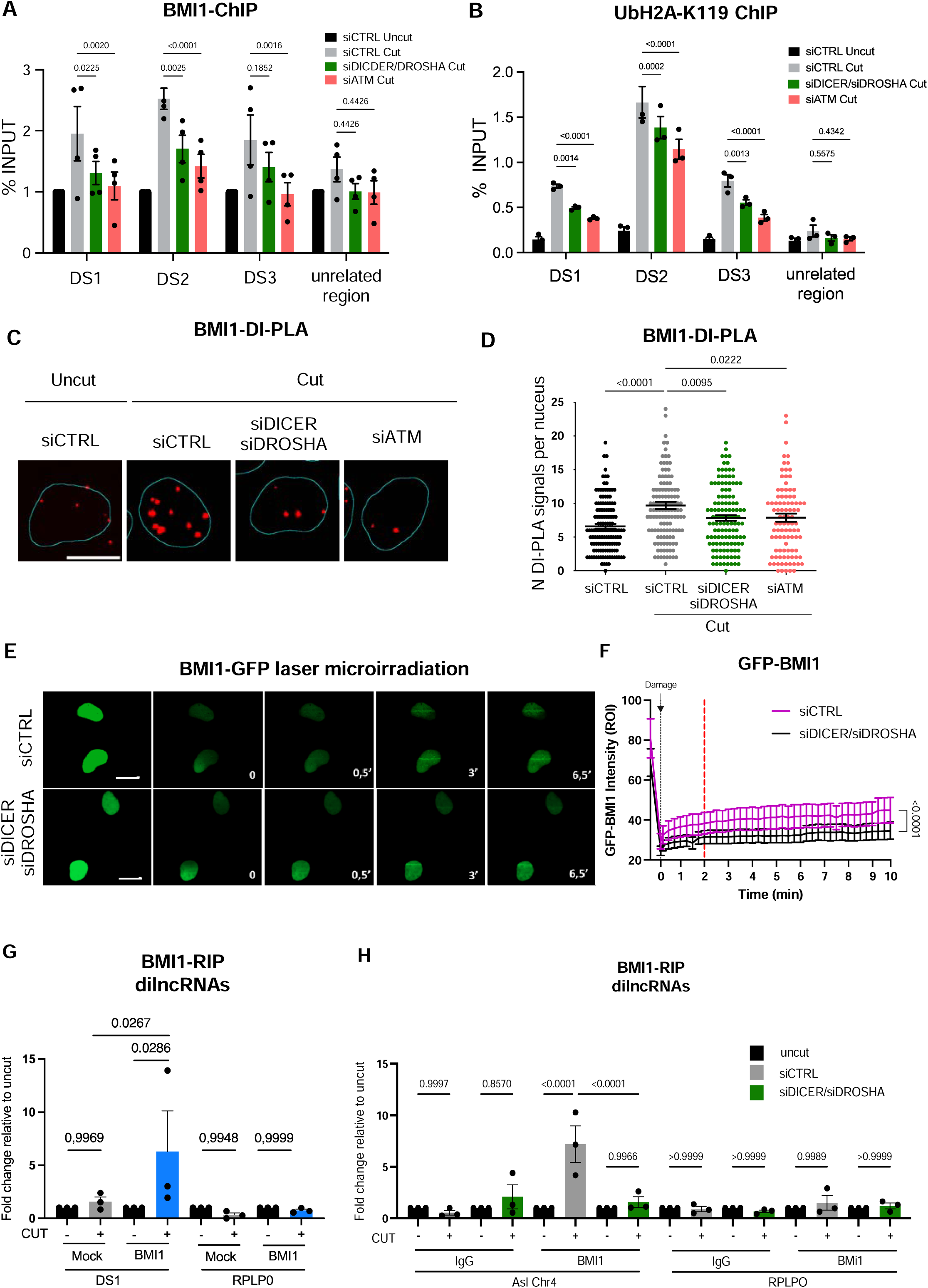
BMI1 is recruited to the damaged site by DROSHA and DICER and interacts with dilncRNA and DDRNAs. **A, B.** ChIP-qPCR analysis at three AsiSI sites (DS1-3) and at an unrelated region, performed in cut and uncut cells treated with non-targeting siRNAs (siCTRL), siRNAs against DROSHA and DICER (siDICER/siDROSHA) or ATM (siATM) transcripts, for BMI1 (A) and ubH2A-K119 (B). DS1 corresponds to an AsiSI break site in the 3’UTR region of RBMXL1 gene; DS2 is located upstream the CDS (promoter region) of CYB561D1 gene; DS3 at 30 bp upstream the 5’UTR region of TRIM37 gene and 500 bp upstream its CDS; the unrelated region corresponds to chr22: 22,373,074; 22,373,287. Error bars represent SEM from three independent experiments. **C.** Representative images of DI-PLA signals for BMI1 recruitment to sites of damage in cut and uncut cells treated with siCTRL, siDICER/siDROSHA or siATM. Scale bar: 20 µm. **D.** Quantification of DI-PLA signals for BMI1 recruitment in cells treated as in C. Error bars represent SEM of more than 150 cells from three independent experiments. **E.** Images of GFP-tagged BMI1 recruitment kinetics to laser micro-irradiation tracks in U2OS cells treated with siCTRL or siDICER/siDROSHA. Scale bars: 20 µm. **F.** Quantification of GFP-BMI1 intensity at laser tracks over time in cells treated as in E. Values are relative to GFP intensity prior to irradiation; error bars represent SEM of at least 12 cells from three independent experiments. Statistical analyses in panels A, B, D and F were performed by One-Way ANOVA. **G,** RNA immunoprecipitation (RIP) of BMI1-associated RNAs followed by RT-qPCR analysis of dilncRNAs generated at the AsISI site on Chr4, performed in uncut and cut DIvA cells. **H,** RIP of BMI1-associated RNAs followed by RT-qPCR analysis of dilncRNA generated at AsISI site on Chr4, performed in uncut and cut DIvA cells treated with siCTRL or DICER/DROSHA co-KD. RPLP0 was used as a negative control. Error bars represent SEM from three independent experiments. Statistical analyses were performed by One-Way ANOVA.

Then, to further demonstrate that DROSHA and DICER control the recruitment of BMI1 to DSB, we employed DNA Damage In Situ Ligation Followed by Proximity Ligation Assay (DI-PLA) ^51^. This technique labels DNA ends generated by a DSB through their ligation to a biotinylated DNA oligonucleotide that is detected by PLA with an anti-biotin antibody. To detect the presence of BMI1 at DNA ends, we combined anti-biotin antibody with anti-BMI1 antibody. We observed that DI-PLA signals for BMI1 increased upon DNA damage generation, again indicating that BMI1 is recruited to DSB DNA ends (Figure 4C,D). Consistent with the result obtained by ChIP-qPCR, DROSHA and DICER co-KD significantly reduced BMI1 DI-PLA signal in damaged cells (Figure 4C,D and S4C). Importantly, BMI1 KD completely abolished BMI1 DI-PLA signal, confirming signal specificity (Figure S4C). Finally, to exclude potential indirect effects, we confirmed by western blotting that BMI1 protein level remained unchanged and that the cell-cycle was unaffected upon DROSHA/DICER co-KD (Figure S4D-F).

The DIvA cell system provides information of a steady-state association of proteins to DSBs, and it is not suitable for monitoring their recruitment kinetics. Thus, to investigate the dynamics of BMI1 recruitment to DNA damage sites and its potential alteration following DROSHA and DICER co-KD, we performed laser micro-irradiation in cells expressing BMI1-GFP. We pre-sensitized U2OS cells with BrdU for 48 hours and then transfected them with BMI1-GFP for 24 hours. Laser micro-irradiation showed an early recruitment of BMI1-GFP to laser stripes that reached plateau around 3-4 minutes post laser-induced damage (Figure 4E,F). As an example, in the same system 53BP1-GFP is recruited to laser stripe later on and reaches plateau at 5-7 minutes post laser (Figure S4G). Notably, DROSHA/DICER co-KD resulted in a reduced accumulation of BMI1-GFP to laser tracks, with a plateau that stably persisted at half of the intensity of control cells despite BMI1 was overexpressed in this settings. These analyses demonstrate that BMI1 recruitment is not only delayed but is constantly dampened in cells inactivated for DROSHA and DICER (Figure 4E,F). Altogether, these results demonstrate that DROSHA and DICER are required for BMI1 recruitment to both individual DSBs and heterogeneous laser-induced DNA damages.

### BMI1 associates with dilncRNAs and ASO or Cas13-mediated targeting of dilncRNAs prevent BMI1 recruitment and DISC

BMI1 has been previously shown to be recruited at specific loci and there repress transcription by interacting with ncRNAs ^52^. Thus, we hypothesized that dilncRNAs may contribute to BMI1 recruitment to DSBs. We performed an RNA immuno-precipitation (RIP) assay by pulling down endogenous BMI1 in cut and uncut DIvA cells and probed for dilncRNA enrichment by strand-specific RT-qPCR at two different AsiSI sites, located on chromosome 4 (Figure 4G) and chromosome 1 (DSB1, Figure S4H). We observed that dilncRNAs generated at both DSB sites were enriched in the BMI1 immunoprecipitated fraction of cut cells (Figure 4G and S4H). *RPLP0* mRNA used as a control showed no enrichment.

Importantly, we also observed that DROSHA co-immunoprecipitated with BMI1 in RIP settings, which preserve RNA integrity (Figure S4I) further confirming the previous result of DROSHA-BMI1 proximity detected by PLA (Figure 3 E-G). Interestingly, BMI1 association with dilncRNAs was strongly reduced in DICER and DROSHA depleted samples (Figure 4H), further suggesting that DROSHA and DICER, or their small RNA products, stabilize BMI1 interaction with dilncRNAs. Thus, we hypothesized that BMI1-dilncRNAs interaction could be supported by DDRNAs via base pairing with dilncRNAs. Therefore, we tested whether BMI1 could also associate with DDRNAs generated upon DSB at repetitive loci. We performed a BMI1 RIP in cut and uncut NIH2/4 cell line ^53^, in which we previously characterized, by small RNA-seq and RT-qPCR, DDRNA accumulation at a specific DSB ^6,17,43^ induced by I-SceI endonuclease ^6,17,43^. In this cell line, the presence of LAC-operon repeats flanking the cut site allows for the visualization of the damaged locus by expressing the LAC-Repressor protein fused to a Cherry-tag. Importantly, we observed that the Bmi1 immunoprecipitated fraction is enriched with DDRNAs with the sequence of the damaged, repetitive site (Figure S4J). Differently, miR-125a and snoRNA U61, two other small non-coding RNAs, were not associated with Bmi1 (Figure S4J). The efficiency of BMI1 IP was evaluated by western blotting in cut and uncut condition (Figure S4K) and the induction of DSB upon I-SceI expression was evaluated by IF as 53BP1- and γH2AX-foci colocalizing with signals of Cherry-LAC, marking the locus targeted by I-SceI endonuclease (Figure S4L). These results indicate that upon DSB formation, BMI1 can specifically associates with DDRNAs, small species more easily detectable in repetitive loci, and dilncRNAs suggesting that both damage-dependent ncRNAs species might be central to mediate BMI1-dependent chromatin modification at DNA damage sites.

We previously used a set of antisense oligonucleotides (ASOs) targeting the dilncRNAs/DDRNA axis to inhibit their function and show their requirement for DDR-foci formation ^6,22,23,54^. To test the impact of dilncRNA/DDRNA inhibition on BMI1 recruitment and DISC, we generated a cellular system in which a DSB can be induced by CRISPR-Cas9 in an integrated GFP reporter gene in HeLa cells (Figure S5A). As already observed in DIvA (Figure 2A,B), we confirmed also in this system that the induction of a DSB within the GFP reporter gene, repress transcription by 50% (Figure 5A,B). Moreover, DROSHA and DICER co-KD, or BMI1 KD, abolish DISC (Figure 5A and S5B; 5B and S5C respectively), without altering GFP expression in uncut conditions (Figure S5D), ruling out indirect miRNA-mediated effects. Similar results were obtained with ATM inhibition (Figure S5E), confirming the role of DICER/DROSHA, BMI1 and ATM in DISC in this additional cellular system in which DSB is induced via CRISPR-Cas9.

**Figure 5:**
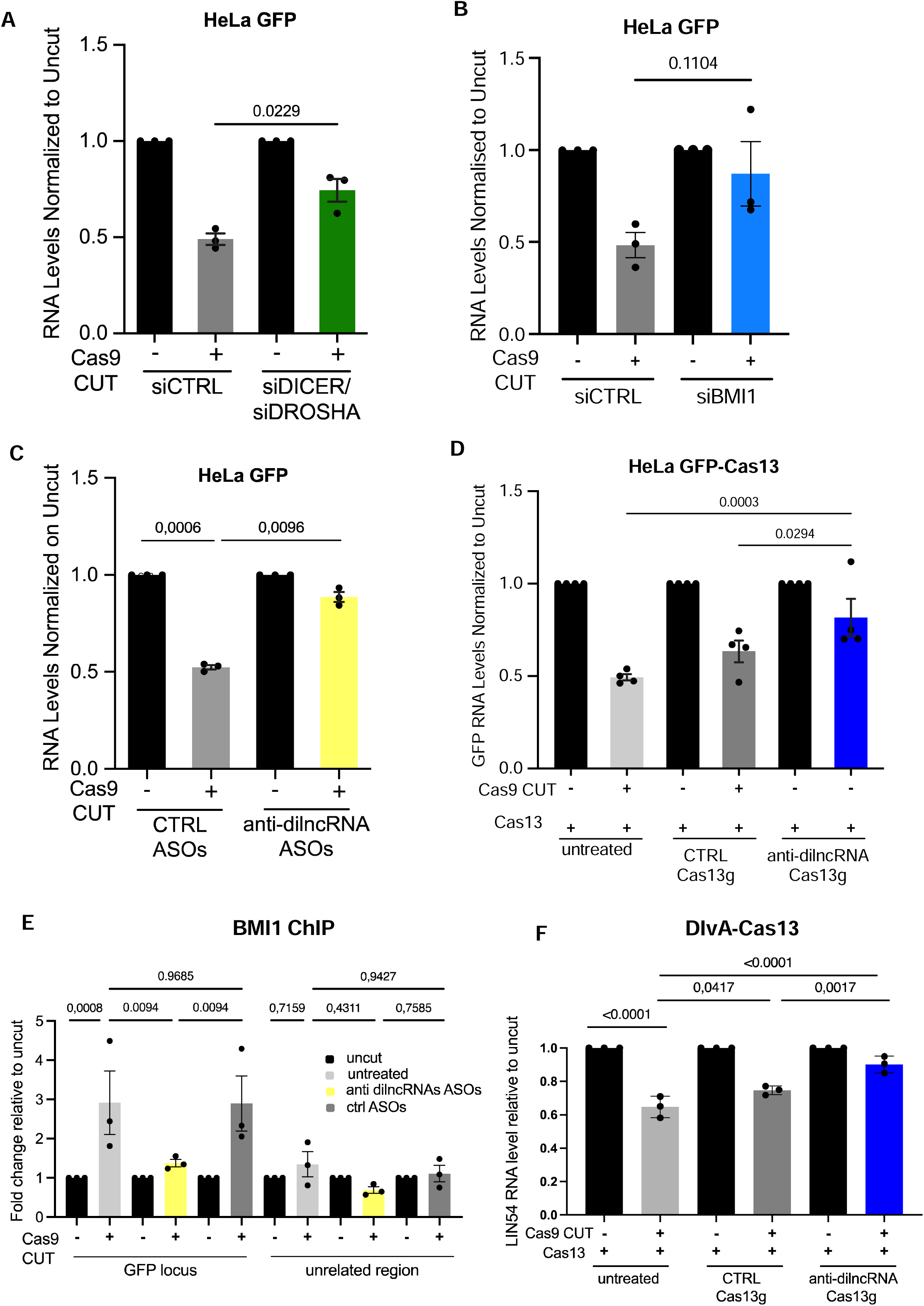
Functional inhibition of the dilncRNA:DDRNA axis by ASOs and its targeting via Cas13d abolishes DISC and BMI1 recruitment to the break. **A.** RT-qPCR analysis of GFP mRNA levels in HeLa cells bearing an integrated GFP gene (HeLa-GFP) treated with control siRNAs (siCTRL) or siRNAs against DROSHA and DICER transcripts (siDROSHA/siDICER). Data are relative to uncut cells for each knockdown condition. Error bars represent SEM from three independent experiments. **B.** RT-qPCR analysis of GFP mRNA levels in cut and uncut HeLa-GFP cells treated with siCTRL or siRNAs against BMI1 transcript (siBMI1). Data are relative to uncut cells for each knockdown condition. Error bars represent SEM from three independent experiments. **C.** RT-qPCR analysis of GFP mRNA levels in cut and uncut cells treated with control ASOs or ASOs targeting dilncRNAs generated at the GFP locus. Error bars represent SEM from three independent experiments. Data are relative to uncut cells for each condition. Statistical test used was paired Student t-test. **D.** RT-qPCR analysis of GFP mRNA levels in cut and uncut HeLa-GFP stably expressing an inducible Cas13d. Cells were treated with a control Cas13 RNA guide (CTRL Cas13g) or dilncRNAs-targeting guides (anti-dilncRNAs Cas13g). Error bars represent SEM from four independent experiments. **E.** ChIP-qPCR analysis for BMI1 at the GFP locus and an unrelated region, performed in uncut (scramble-guide-Cas9) and cut (GFP-guide-Cas9) HeLa-GFP cells treated with control ASOs or dilncRNAs-targeting ASOs. Data are relative to uncut cells after the subtraction of mock values for each treated condition. Error bars represent SEM from three independent experiments. **F.** RT-qPCR analysis of LIN54 (DSB at Chr4) mRNA levels in cut and uncut DIvA cells stably expressing an inducible Cas13d. Cells were treated with a control Cas13 RNA guide (CTRL Cas13g) or guides targeting dilncRNAs generated downstream the AsI cut site on Chr4 (Chr4 RNA Cas13g). Error bars represent SEM from three independent experiments. Statistical analyses in panels B, D, E and F were performed by One-Way ANOVA.

Then, to directly probe dilncRNA role in DISC, we transfected ASOs targeting dilncRNAs generated at the DSB (Figure S5A, Table3), along with plasmids encoding Cas9 and a guide RNA (either scrambled or directed to the GFP locus), and measured the relative GFP expression by RT-qPCR. Strikingly, the administration of ASOs targeting dilncRNAs prevent DISC, while control ASOs do not (Figure 5C). Importantly, ASO transfection did not enhance *per se* GFP expression in undamaged cells, while, as expected, anti-GFP ASOs reduced it significantly (Figure S5F).

To independently support the notion that the dilncRNAs:DDRNAs axis control DISC, we used the recently identified Cas13d variant^55^ as an additional approach to specifically target dilncRNAs ^56^. We therefore inserted in the GFP reporter cell system a doxycicline inducible form of the Cas13d enzyme and transfected multiple plasmids expressing five Cas13d guide RNAs targeting dilncRNA generated at CRISPR-Cas9 DSB (anti-dilncRNAs Cas13g), or a control gRNA (CTRL Cas13g), targeting an unrelated region (Figure S5A and Table 4). Strikingly, we observed that the expression of Cas13d gRNAs targeting dilncRNAs at the GFP locus significantly reduced DISC, while the control gRNA did not significantly impact on it (Figure 5D). These results demonstrate that dilncRNAs play a direct role in DISC.

Since we showed that dilncRNAs promote DISC and interact with BMI1 (Figure 4 and 5A-D), we tested if their inactivation using ASOs could interfere with BMI1 recruitment to DSB. We therefore performed a ChIP-qPCR for BMI1, γH2AX and phospho-ATM upon CRISPR-Cas9 cut, in cells transfected with ASOs targeting dilncRNAs or control one. We observed that BMI1 was enriched at a γH2AX- and pATM-positive DSB generated by CRISPR-Cas9 in the GFP locus, but not in an unrelated region of the genome (Figure 5E and S5G,H). Strikingly, ASOs targeting dilncRNAs, but not control ones, abolished BMI1 recruitment, while leaving γH2AX and pATM levels unaltered (Figure 5E and S5G,H). These results further support a model in which dilncRNAs generated at the site of DNA damage, together with DROSHA and DICER, control DISC by recruiting BMI1 to repress transcription of adjacent genes.

Finally, to confirm that dilncRNAs also control DISC at endogenous loci in the DIvA cellular system, we used the Cas13d approach to target the dilncRNA induced at the DSB on Chr4 and evaluate the expression of LIN54 (Figure S5I), the endogenous protein-coding gene located a few hundred bases upstream to the DSB. For this purpose, we infected DIvA cells with the same doxycycline inducible form of Cas13d enzyme, and then transfected cells with either dilncRNA-targeting Cas13d guides or the same amount of scrambled control Cas13d guide (Table 4). Firstly, we could confirm that LIN54 is repressed when the DSB is induced in its proximity, and this event was dependent on DICER/DROSHA (Figure S5I). Importantly, we could show that the transfection of Cas13d guides targeting dilncRNAs generated on one side of the break respect to the gene unite, was sufficient to strongly dampen DISC of LIN54, while scramble control guide did not (Figure 5F). Moreover, expression of Cas13d alone had no effect on DISC (Figure 5F). As additional controls, we could show that the transfection of dilncRNA-targeting Cas13d guides in the absence of Cas13d expression, had no impact on DISC of LIN54 (Figure S5J), demonstrating that loss of DISC is mediated by Cas13d-dependent dilncRNA cleavage. Moreover, using a mix of non-targeting guides (Table 4), among which we included the GFP targeting guide, which was dampening DISC in the HeLa-GFP system, (Table 4), we did not observe any impact on DISC (Figure S5J), highlighting the specificity of Cas13d system in our two different experimental settings.

In summary, this study provides evidence that *de novo* synthesis and processing of ncRNAs induced by DSBs are not only required for DDR activation but also control DISC at the same loci.

## Discussion

It is now well established that DSBs, generated by different means, cause transcriptional repression of neighboring genes. The dampening of active transcription in the vicinity of the damage, a phenomenon recently named by others with the acronym DISC ^33^, is not just a passive event caused by the physical interruption of the DNA template. Rather, it is an active process that involves the recruitment of the repressive complex PRC1 to damaged chromatin in an ATM-dependent manner. Concomitantly, *de novo* transcription starting from DNA ends has been now consistently observed in different studies and contexts ^18,57^. Indeed, we previously reported that dilncRNAs can be generated at the site of damage both in exogenously integrated repetitive loci ^17^ and at endogenous loci, including deprotected telomeres ^22^ and CRISPR/Cas9-mediated cleavages ^6^.

In this work, for the first time, we mechanistically connect the two seemingly contradictory events, DISC and damage-induced *de novo* transcription, showing how endogenous non-coding RNAs, together with the RNA interference (RNAi) machinery, control chromatin state and consequently transcription of protein coding genes of the same locus (Figure 6A). This may be functionally and perhaps mechanistically related to what was described in *S. pombe* for depositing, and then limiting, constitutive heterochromatin at centromeres ^58–61^. More in general, the role of RNAi factors in chromatin-mediated transcriptional control is extensively reported ^62–65^.

**Figure 6:**
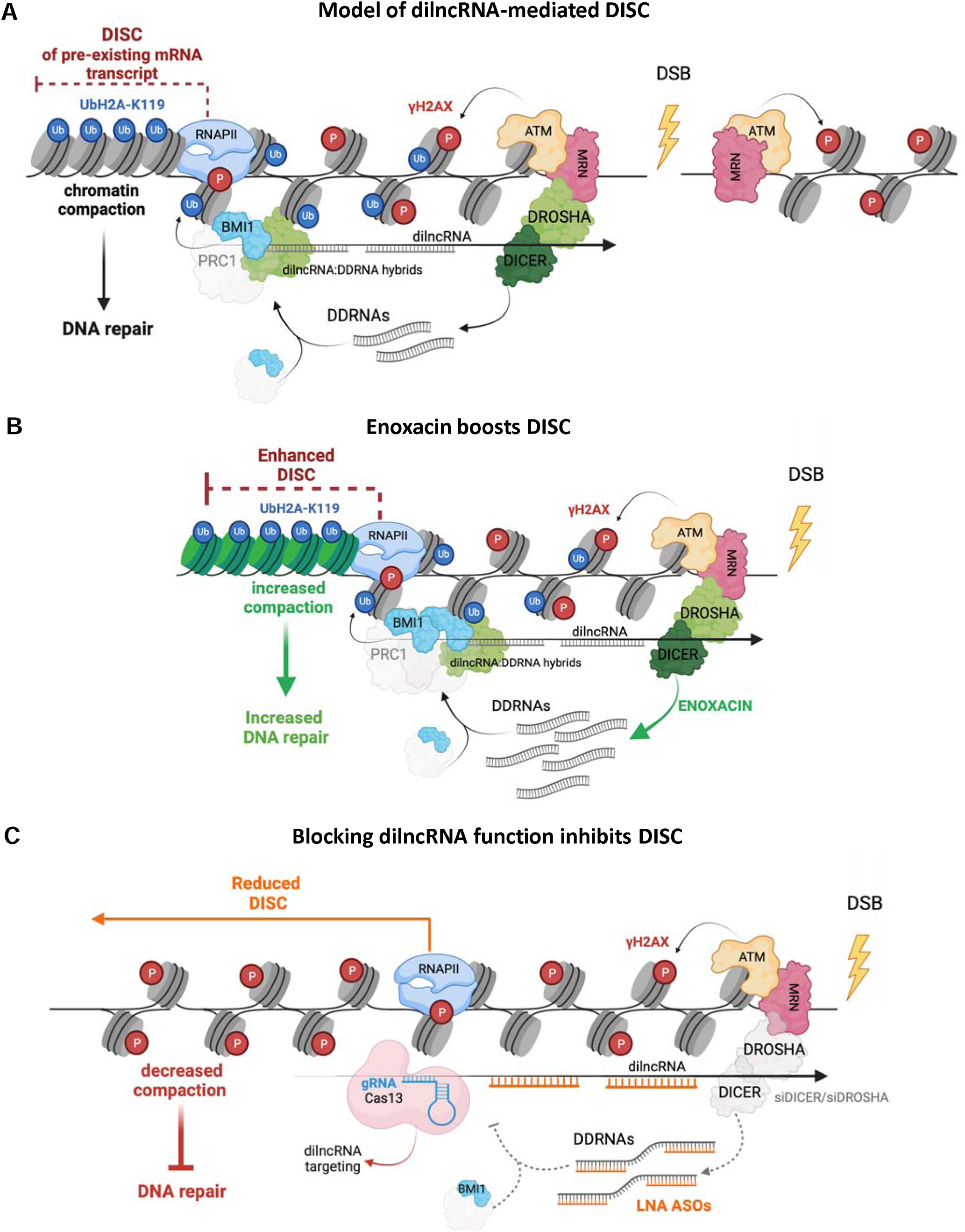
Schematic representation of the proposed model. **A.** Model of DISC mediated by the dilncRNAs:DDRNAs axis. In physiological conditions, when a DSB is generated dilncRNAs are *de novo* transcribed and processed by DROSHA and DICER into DDRNAs, which sequence-pair with dilncRNAs, recruiting the PRC1 complex at the chromatin. Here, BMI1 deposits the UbH2A-K119 histone mark, which leads to chromatin compaction and silencing of the pre-existing mRNA transcript (DISC), thus facilitating proper DNA repair. **B.** Enoxacin boosts DISC. When enoxacin is administered, DICER processing activity is stimulated and the increased DDRNA production leads to an increased chromatin compaction, enhancing DISC. **C.** Blocking DDRNA function inhibits DISC. DICER/DROSHA depletion, ASOs-dependent inhibition or Cas13d-mediated dilncRNA targeting abolishes DISC. The figure was created with BioRender.com.

We show that the repression of canonical genes upon DSBs requires DROSHA, DICER and dilncRNAs, which allow for the deposition of the Ub-H2AK119 repressive marker by recruiting to the damaged site the ubiquitin ligase BMI1, subunit of the PRC1 complex. This provides new mechanistic details of how a damaged locus mediates PRC1 recruitment, chromatin modification and DISC in a sequence-specific fashion. Other published reports support the notion that DROSHA, DICER and their small RNA products control the recruitment to DSBs of chromatin modifiers, such as the methyltransferase MMSET and the acetyltransferase Tip60, resulting in a local flexible chromatin configuration that allows the recruitment of key repair factors and thus promotes DSB repair ^4^.

Enoxacin effects can be used as a read out of DICER functioning in the cell. By reporting its ability to stimulate DISC, we prove that this mechanism is specifically dependent on DICER endoribonuclease activity, and thus very likely on DDRNA production. Beyond the mechanistic detail, this is of relevance because it highlights how a single small molecule can amplify different aspects of DDR signaling (Figure 6B), resulting in an enhanced DNA damage repair, too ^43^. This result also strengthens the notion that DISC and repair are intimately connected and could pave the way for a potential broader applicability of the use of Enoxacin molecule in reinforcing physiological repair to contrast pathological events.

Multiple mechanisms converge on ATM activation to repress canonical transcription of DSB-flanking genes ^66,67^. For instance, ATM has been recently shown to recruit the Bromodomain Containing 7 (BRD7) protein to damaged chromatin, bringing the PRC2 and NuRD complexes together to achieve DISC ^67^. Additionally, ATM recruits via MDC1 the E3 ubiquitin ligases RNF8 and RNF168, which both have a role in repressing transcription by promoting RNAPII pausing or chromatin compaction ^30,68,69^. Finally, different ATM substrates are involved in transcriptional silencing at the breaks, including the BAF180 subunit of the PBAF remodeling complex, required for Ub-H2AK119 deposition ^26^, and the Ser/Thr kinase DYRK1B ^33^, which recruits PBAF components to damaged chromatin ^33^. Though, by using Enoxacin we demonstrate here that DICER’s nuclease activity can contribute to DISC also separately from ATM, by showing that ATM’s kinase activity is dispensable for enoxacin-mediated DISC stimulation. This is also in line with the fact that ATM inhibition does not affect DROSHA recruitment at the break site ^13^ nor dilncRNA transcription ^6^. Differently, not only we observe that MRN inhibition alone is sufficient to abolish DISC, but also that MRN activity is necessary for DISC stimulation by Enoxacin. This is indeed coherent with the observation that MRN’s activity is necessary for both dilncRNA transcription and DROSHA recruiting at the damage site ^13^. These results indicate that MRN might function in DISC also independently from its role in ATM activation at DNA damage site. Indeed, some reports in the last year suggested that MRN have important roles in DDR signaling and repair which are independent from ATM ^70–72^. Our evidence that Enoxacin can stimulate DISC only if DROSHA is recruited to DSB, but independently from ATM places dilncRNA/DDRNA axis in a parallel, if not upstream, position respect to ATM activity in the DSB signaling cascade.

Importantly, a series of parallel pathways, along with ATM-dependent events, has been already described to coordinate the transcriptional silencing around damaged genes in an ATM-independent fashion, such as the DNA-PK-promoted RNAPII pausing and degradation ^25,73^ or the PARP1-dependant chromatin modifications ^31,74^. In our work, we propose an additional mechanism whereby dilncRNA/DDRNA production coordinates BMI1 recruitment and chromatin modification to achieve DISC, independently from the ATM kinase activity.

Intriguingly, here we report an interaction between DROSHA and BMI1 which is RNA dependent, and we show that indeed both dilncRNAs and DDRNAs associate to BMI1. One possible model that one can envision, also supported by other previous publication ^6^, is that BMI1 associate with DDRNAs which in turn anneals with dilncRNAs. Indeed we could show that BMI1 association with dilncRNA is strongly reduced in cells knocked down for DROSHA and DICER, thus devoid of DDRNAs. On the other side, we also observed that DROSHA protein itself associate with BMI1 via RNA. It was already demonstrated that BMI1 interacts with the MRN complex ^48^ and so DROSHA does ^13^. In addition, dilncRNA transcription is MRN complex dependent ^11^. We speculate that a first interaction of BMI1with the MRN complex and the dilncRNAs may promote its early recruitment, while its subsequent interaction with DROSHA and DDRNAs could support its retention at the break, two aspects necessary for rapid and sustained gene silencing, respectively. Indeed, BMI1 is an early responder, as confirmed by our live imaging results, but it can be visualized at the site of damage at much later time points as well ^75^. Importantly, the interaction with dilncRNA:DDRNA pairs is pivotal for BMI1 recruitment at the break, since administration of sequence-specific ASOs complementary to dilncRNAs completely abolishes BMI1 localization at the damage site and DISC (Figure 6C) and DICER/DROSHA co-KD reduces the amount of dilncRNAs associated to BMI1. Noteworthy, mounting evidence shows that long and short non-coding RNAs guide the repressive PRC1/2 complexes towards their target genes, ensuring specific chromatin modification patterns and transcriptional landscapes, both in terms of constitutive and facultative heterochromatin ^52,76–78^. This is line with a role of dilncRNAs in directing a sequence-specific recruitment of the BMI1/PRC1 complex to the site of break, where it can exert its repressive function and support repair. Since, BMI1 was reported to interact with MRN complex upon DSB induction ^50^ and we demonstrated that DROSHA binds to MRN ^13^ we entertain the possibility that MRN mediates the early recruitment to DSB of both BMI1 and DROSHA, a model consistent with our observation that DISC is exquisitely MRN-dependent.

Given the crucial role of DROSHA and DICER canonical miRNA products in regulating transcript stability and translation, we aimed to exclude any their indirect effect in DISC. Of note, miRNAs have been shown to regulate the expression of different DNA repair factors ^79^, while others are found to target the progression of cell cycle ^80,81^, thus indirectly influencing DNA repair. Nevertheless, miRNAs have not been linked to transcriptional silencing of damaged genes, and indeed we could show that they do not affect DISC in our system. First, we show that cell cycle progression is not affected by either Enoxacin administration or DROSHA and DICER silencing, ruling out indirect effects on DNA damage repair. Moreover, we prove that inactivation of GW182-like proteins, RNA-interference components required for miRNA-mediated translational repression ^82^, does not affect DISC. Similarly, inhibition of the miRNA-guided gene silencing via the expression of the T6B peptide ^41^ does not impact on DISC (Figure S2F), demonstrating that DROSHA and DICER act in DISC independently from their role in miRNA biogenesis.

Previous studies proposed that transcriptional downregulation is achieved by exclusion of RNAPII from damaged chromatin ^83^. On the other hand, other reports suggest that occupancy of total RNAPII at damaged chromatin is not affected ^24^, while only the amount of the elongating form of RNAPII (Ser2-phosphorylated) is reduced upon damage generation ^24^. Indeed, in the DIvA system, we previously observed that RNAPII elongation is reduced in the chromatin surrounding the DNA lesion, while the total amount of RNAPII protein remains mostly unaltered ^29^. This evidence may suggest the existence of a controlled switch in RNAPII elongation. It is tempting to speculate that dilncRNA/DDRNAs and PRC1 complex induce a rapid switch from transcription elongation to RNAPII pausing, without the need of totally removing RNAPII from damaged chromatin.

Our data obtained by laser micro-irradiation shows that global transcription decreases after DNA damage, underlining how dilncRNAs are low-abundance transcripts mostly kept chromatin bound. This is in agreement with a model in which local dilncRNA transcription and DDRNA production are not sufficiently abundant to directly induce the immediate degradation of gene transcripts coming from the same locus by anti-sense annealing. On the contrary, there is a need to recruit a chromatin modifier that modulates chromatin ubiquitination and possibly compaction to dampen transcription of DSB-flanking genes. Consistently, in such a model, DSBs and the newly generated dilncRNA recruit the PRC1 complex to modulate chromatin state and promote the switch of the RNAPII machinery from an active, processive state to a pausing one. Indeed, the loss of DROSHA and DICER restores transcription in damaged, γH2AX positive chromatin. Overall, as already described for similar mechanisms such as X-chromosome inactivation ^84^ or the deposition of centromeric constitutive heterochromatin ^85,86^, the emerging model suggests that the synthesis of ncRNAs together with factors of the RNAi machinery is required for the transcriptional repression of DSB-flanking genes requires (Figure 6A-C).

## Methods

### Cell culture

U2OS cell line was cultured in DMEM (Euroclone; #ECB4004L) supplemented with 10% FBS, 1% L-glutamine and 1% P/S. U2OS 2-6-3 cells were cultured as already described ^24^. To induce transcription of the reporter locus, 1 μg/ml Doxycycline was added to complete growth medium for 3 hours before stopping the experiment. U2OS MDC1–GFP and U2OS 53BP1–GFP cells were grown as shown before (Hable et al., 2012) ^20^. DIvA (AsiSI-ER-U2OS) cell line was cultured as done previously ^38^. DNA damage in DIvA cells was induced with 300 nM 4-OHT (Sigma-Aldrich; #H7904) for 4 hours. When indicated, DIvA cells were treated with 25 µM Enoxacin (Sigma-Aldrich; #557305-250mg) dissolved in water and added to the growth medium for 48 hours; 100 µM MRN inhibitor Mirin (Sigma; #M9948) dissolved in DMSO and added to the growth medium for 4 hours; 5 µM ATM inhibitor KU-60019 (Selleckchem; #S1570) dissolved in DMSO and added to the growth medium overnight.NIH2/4 cells were grown as already described ^87^. To induce DNA damage, NIH2/4 cells were transduced with a lentiviral I-SceI-GR expressing construct or with an empty vector as a control ^6^. I-SceI-GR was translocated to the nucleus by treating cells with 0.1 µM triamcinolone acetonide (Merck; #T6501) for 1.5 h before harvesting.

We generated HeLa cells with an integrated GFP locus (HeLa-GFP) by transducing cells with the pLenti-CMV-MCS-GFP-SV-puro lentiviral plasmid (Addgene Plasmid #73582). Cells were cultured in DMEM (Euroclone; #ECB4004L) supplemented with 10% FBS, 1% L-glutamine, 1% P/S. To select the GFP construct, transduced HeLa cells were kept under selection by adding 1 μg/ml Puromycin to the culture medium.

A doxycycline-inducible Cas13d HeLa-GFP cell line was then generated by transducing HeLa-GFP cells with the inducible RfxCas13d encoding pLentiRNACRISPR_007-TetO-NLS-RfxCas13d-NLS-WPRE-EFS-rtTA3-2A-Blast lentivirus (Addgene Plasmid #138149). Transduced cells were maintained in DMEM (Euroclone; #ECB4004L) supplemented with 10% FBS, 1% L-glutamine, 1% P/S, and selected by adding 1 μg/ ml Puromycin and 2 μg/ ml Blasticidin to the culture medium. All cells were grown at 37°C in 5% CO_2_.

### RT-qPCR

Standard and strand-specific RT-qPCR was performed as described in ^6^. Briefly, total RNA was extracted using the RNEasy Mini Kit (Qiagen; #74106) according to the manufacturer’s instructions. The isolated RNA was then treated with TURBO Dnase (Invitrogen; #AM1907) according to the manufacturer’s instructions. For standard RT– qPCR, 1 µg of RNA was reverse-transcribed using the SuperScript^TM^ IV First-Strand Synthesis System Kit (Invitrogen; #18091050) by using random examers. For strand specific RT-qPCR, 500 ng of RNA were reverse-transcribed using the SuperScript^TM^ III First-Strand Synthesis System Kit (Invitrogen; # 18080051) by using gene-specific and strand-specific primers (DSB1 FW and RPLP0 REV for dilncRNA and negative control detection, respectively). Real-time quantitative PCR (qPCR) reactions were performed on a LightCycler® 480 II Sequence Detection System (Roche) by using QuantiTect SYBR® Green PCR kit (Qiagen; #204145). For a complete list of the used primer sequences see Table 1.

**Table 1:**
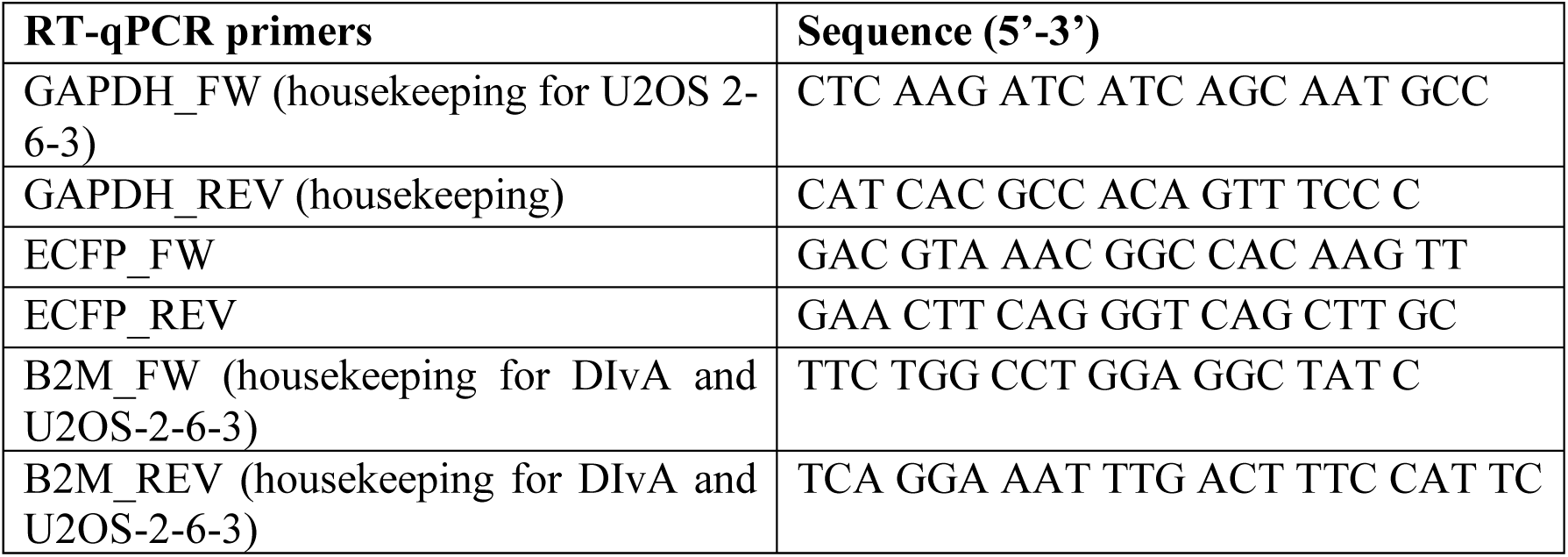
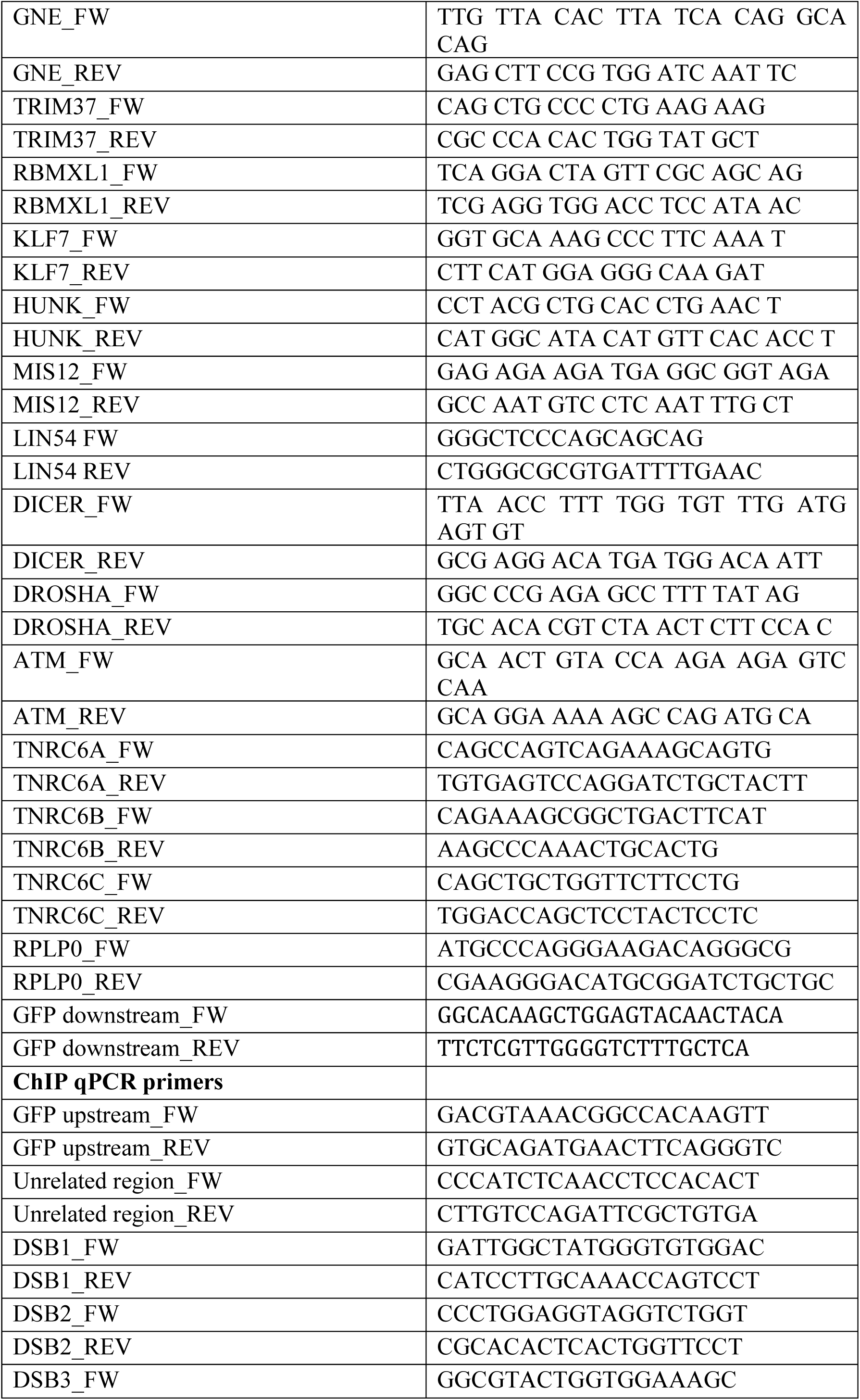
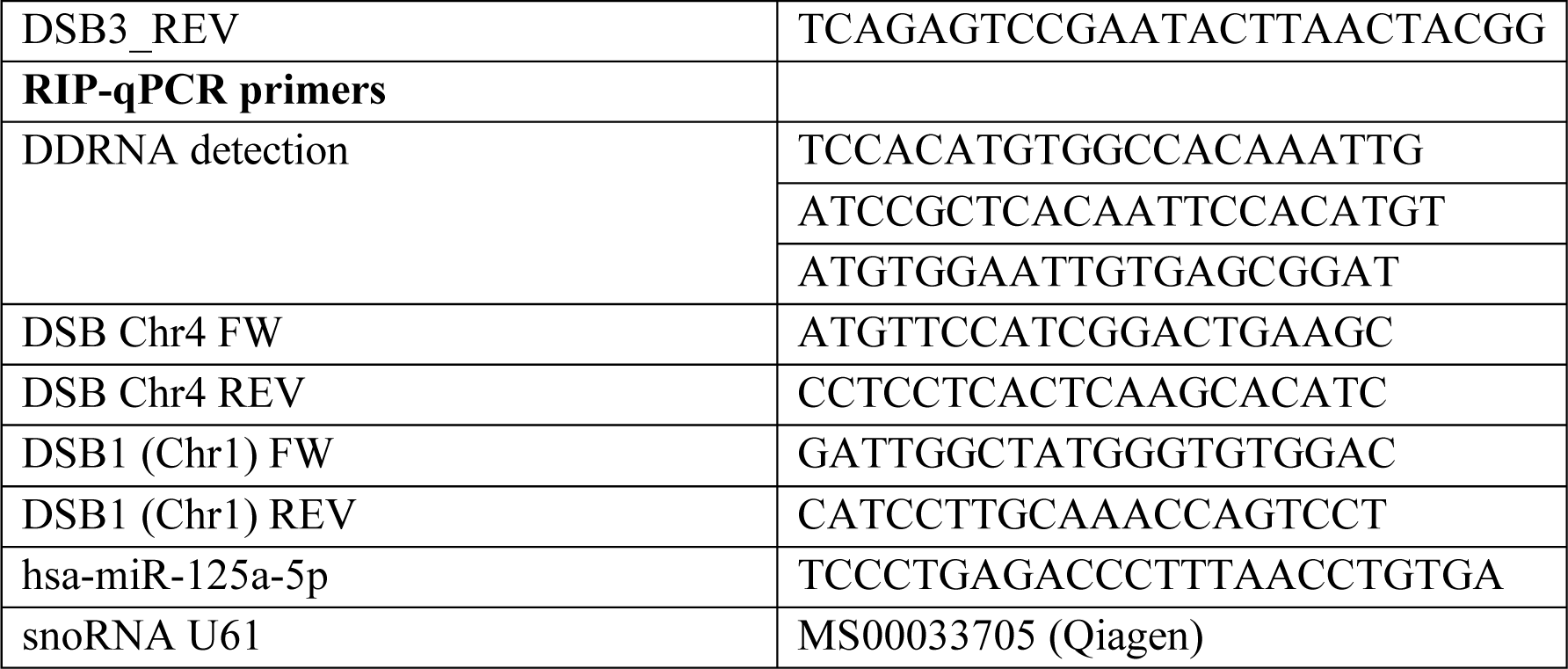
primers used in this study.

### Transfections

For RNA interference, cells were transfected with 5-20 nM siRNA oligos (Table 2) using Lipofecatmine RNAiMAX transfecting agent (Invitrogen; #13778-075) according to the manufacturer’s instructions for 72 hours. mCherry-LacI-FokI-WT and mCherry-LacI-FokI-D450A expressing vectors ^24^ were transfected in U2OS 2-6-3 cells using Lipofectamine LTX (Invitrogen; #15338-100) for 24 hours. T6B wild type (WT) and T6B inactive mutant expressing vectors ^41^ were transfected in DIvA cells with Lipofectamine 2000 (Invitrogen; #11668-027) for 24 hours before RNA extraction. Cherry-LacR expressing vector ^53^ was transfected with Lipofectamine 2000 (Invitrogen; #11668-027) in NIH2/4 cells as previously described ^6^. For laser micro-irradiation experiments, BMI1-GFP expressing vector (Addgene Plasmid #128328) was transfected in U2OS cells by Lipofectamine 2000 (Invitrogen; #11668-027) 24 hours before cell irradiation. For experiments in HeLa-GFP cells, Cas9 expressing vectors (Addgene) bearing either a non-targeting scrambled gRNA or a cut-inducing GFP-targeting guide gRNA (see Table 4 for sequences of scramble gRNA and GFP-targeting gRNA) were transfected using Lipofectamine 2000 (Invitrogen; #11668-027) for 24 hours before cell fixation or RNA extraction. Where indicated, a pool of antisense oligonucleotides (ASOs, RA1-A5; B2-B5) was concomitantly transfected at a final concentration of 50 nM. For a complete list of the ASOs sequences used in this study see Table 3. For experiments in inducible Cas13d HeLa-GFP, cells were transfected with the Cas9 plasmids and Cas13 plasmids (1:2 ratio) using Lipofectamine 2000 (Invitrogen; #11668-027) for 24 hours before RNA extraction. Cas13 expression was induced by adding 1 µg/ml Doxycycline to the culture medium for at least 16 hours before RNA extraction.

**Table 2:**
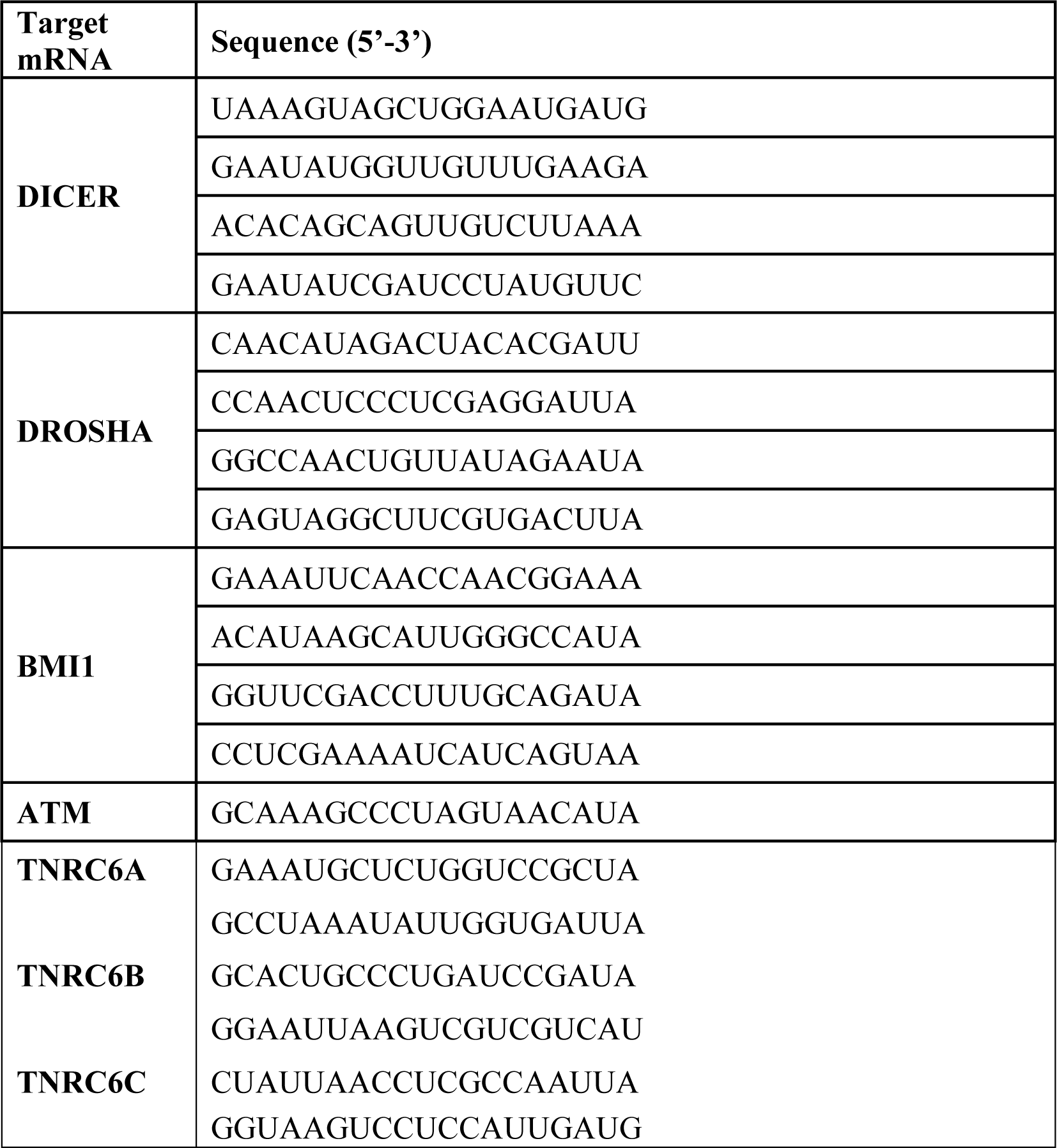
siRNAs used in this study.

**Table 3:**
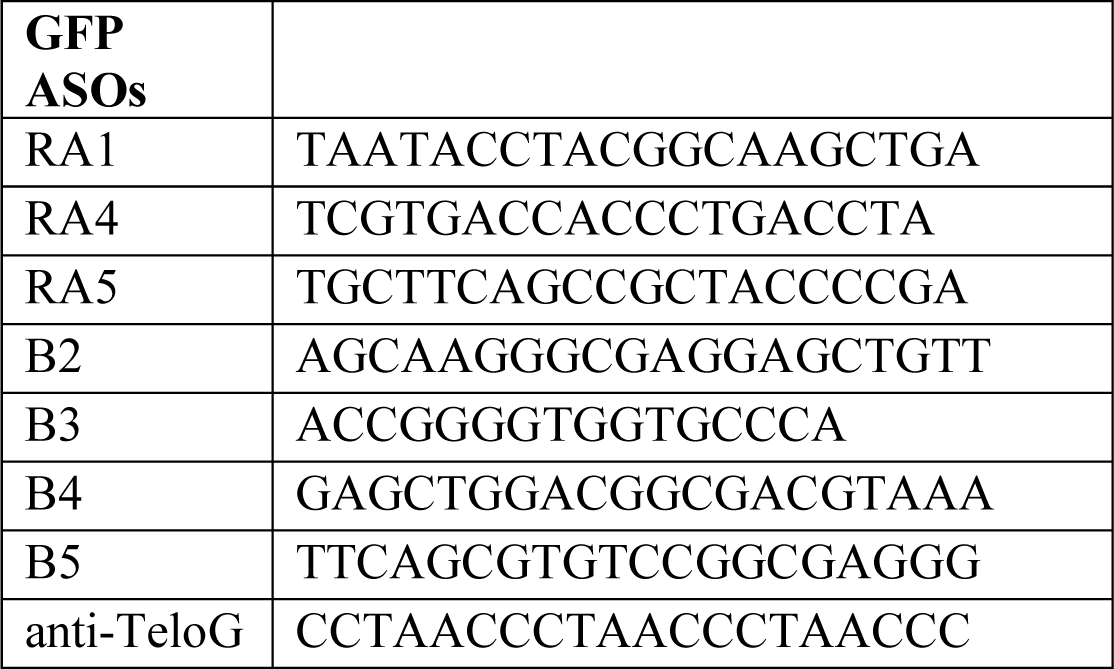
ASOs used in this study.

### Laser micro-irradiation

Laser-induced DNA damage in U2OS MDC1-GFP was performed as described in Francia S. et al. JCS 2016 ^20^, using a Leica TCS SP5 point scanning confocal microscope equipped with a Leica HCX PL APO 63×/1.4NA oil immersion objective and an environmental microscope incubator (OKOLab) set to 37°C and 5% CO_2_ perfusion. The Leica TCS SP5 confocal microscope was driven by Leica LAS AF software. Laser micro-irradiation was carried out using a 50 mW 405 nm diode laser with a 100% power output. At 2× digital magnification, multiple regions of interest (ROI) of the same size were selected in each nucleus and the 405 nm laser was used to scan the ROIs for 50 iterations (dwell time per pixel 0.49 μs).

Both γH2AX and EU signal intensity was quantified with CellProfiler software ^88^ masking on γH2AX stripes. The EU intensity of individual stripes was normalized for the γH2AX signal in the same area, as a reference of DNA damage induced by the laser. Laser-induced DNA damage for GFP-BMI1 transfected U2OS cells and GFP-53BP1 U2OS cells was performed using a Zeiss LSM800 confocal linear laser-scanning microscope (Zeiss LSM 800) equipped with four lasers [diode laser 405 nm (5 mW), diode laser 488 nm (10 mW), diode laser 561 nm (10 mW) and diode laser 640 nm (5 mW)], two Master gain with high sensitivity and a 63×/1.4NA objective and an environmental Incubator Unit XL multi S1 (Pecon) set to 37°C and 5% CO_2_ perfusion. The ZEISS LSM 800 was driven by Zeiss Zen Blue 2.6 software. Laser micro-irradiation was carried out using 5mW 405 nm diode laser with a 100% power output. At 0.5× digital magnification, multiple regions of interest (ROI) of the same size were selected in each nucleus and the 405 nm laser was used to scan the ROIs for 400 iterations (dwell time per pixel 0.44 μs).

In both cases, cells were cultured in glass-bottom dishes and pre-sensitized for 72 hours by adding 10 μM BrdU to culture medium.

### 5-ethynyl uridine (5-EU) incorporation

5-ethynyl uridine (5-EU) staining was performed as described in ^45^. Following DNA damage generation via laser micro-irradiation, nascent RNAs were labelled with 1 hour pulse of 1 mM 5-EU. Cells were then fixed in 4% paraformaldehyde (PFA; ChemCruz; #sc-281692) and levels of EU incorporation was detected by Click-iT RNA imaging kit (Invitrogen; #C10330) according to the manufacturer’s instructions.

### Indirect Immunofluorescence (IF) and imaging analysis

Immunofluorescence for DDR markers was performed as described in ^17^. For experiments in U2OS 2-6-3 reporter cells, more accurate representative images from one experiment were acquired with a confocal linear laser-scanning microscope (Zeiss LSM 800). For experiments in laser micro-irradiated DIvA cells, representative images from one experiment were acquired with a confocal laser microscope (Leica TCS SP2) by sequential scanning. Comparative immunofluorescence analyses were performed in parallel with identical acquisition parameters and exposure times among conditions, using CellProfiler software ^88^ for experiments in DIvA cells and HeLa-GFP cells; ImageJ Software (NIH) was used for experiments in U2OS 2-6-3 rt-TA YFP-MS2 cells and laser micro-irradiated DIvA cells; ZEN Blue Software (Zeiss) was used for experiments in GFP-BMI1 transfected U2OS cells. For the complete list of antibodies used in this study see Table 5.

### RNA-sequencing and data analysis

For total RNA-seq in DIvA cells, total RNA was isolated using the RNeasy Kit (Qiagen; #74106) according to the manufacturer’s instructions. mRNA-seq indexed library preparation was performed starting from total mRNA (Illumina, TruSeq Stranded mRNA) according to the manufacturer’s instructions. Library quality and quantity was assessed on the 2100 Bioanalyzer High Sensitivity DNA kit (Agilent; #5067-4626), quantified on Qubit dsDNA HS Assay, normalized and pooled to perform a multiplexed sequencing run. Clusters were generated on the Illumina flow cell and sequencing was carried out on NextSeq 550 System with paired-end 75 bp. All six conditions were sequenced as biological triplicates. The reads for RNA-seq experiments were aligned to the GRCh37/hg19 assembly human reference genome using the nf-core RNA-seq pipeline (https://nf-co.re/rnaseq); in particular the alignment was performed with the STAR aligner ^89^ and quantification was performed with Salmon (https://combine-lab.github.io/salmon/). Differential gene expression analysis was performed using the Bioconductor package DESeq2 (Love et al., 2014) that estimates variance-mean dependence in count data from high-throughput sequencing data and tests for differential expression exploiting a negative binomial distribution-based model. All RNA -sequencing data have been loaded on GEO (Submission GSE193821).

### Western Blotting

Immunoblotting was performed ad described in Cabrini M. et al JCS 2021 ^13^. Proteins were visualized by chemiluminescence detection using Luminata Classico or Crescendo (Millipore) followed by autoradiography on ECL films (Amersham), or using ChemiDoc^TM^ MP Imaging System (Biorad). For the complete list of antibodies used in this study see Table 5.

### Chromatin immunoprecipitation (ChIP)

ChIP in DIvA cells was performed as already described ^13^. ChIP in HeLa cells was conducted as shown previously ^90^. Immunoprecipitation was carried out overnight at 4°C with magnetic beads (LifeTechnologies; #10003D) previously bound to specific antibodies or IgG for mock. DNA was cleaned up by QIAquick PCR purification column (Qiagen; #28506) according to the manufacturer’s instructions, eluted in 30 µl of elution buffer and analysed by SYBR® Green PCR kit (Qiagen; #204145). In DIvA cells DS1, 2 and 3 correspond to AsiSI-63, AsiSI-507 and AsiSI-453 respectively. Unrelated region falls in chromosome 22. For the complete list of antibodies used in this study see Table 5.

### Proximity ligation assay (PLA)

To perform the PLA assay, cells were fixed with 4% paraformaldehyde for 10 minutes and permeabilized with 0.5% Triton X-100 for 10 minutes at room temperature. Coverslip were blocked with the blocking reagent provided in the Duolink Kit (SIGMA; # DUO92002) for 30 minutes at 37° C and then incubated with the indicated primary antibodies for 1 hour at room temperature. After incubation with primary antibodies, cells were washed with PLA Buffer A (SIGMA; #DUO82049) and incubated with PLUS (SIGMA; #DUO92002) and MINUS (SIGMA; #DUO92004) probes for 1h at 37° C. Then, ligation and amplification steps were performed according to the manufacturing instructions of the Duolink In Situ Detection Kit (SIGMA; #DUO92007). At this point, cells were washed twice with PLA Buffer B (SIGMA; #DUO82049) for 5 minutes at room temperature and once with Buffer B 0.01× (SIGMA; #DUO82049) for 1 minute at room temperature. Finally, covers were stained with DAPI for 2 minutes at room temperature and mounted for microscope acquisition and subsequent analysis.

### DNA damage in situ ligation proximity ligation assay (DI-PLA)

DI-PLA was performed as previously described in ^13,91^. Briefly, cells were fixed and permeabilized as described above in the PLA section. Coverslips were then washed twice for 5 min in 1x rCutSmart buffer (NEB; #B6004S) and once in 1x blunting buffer (NEB; # B1201SVIAL). Afterwards, blunting was performed at room temperature for 60 min in a final volume of 50 μl for each coverslip [38.5 μl H 2O, 5 μl 10× blunting buffer (NEB; # B1201SVIAL), 5 μl dNTP 1 mM (NEB; # N1201AVIAL), 0.5 μl BSA (20 mg/ml; NEB; #B9200S) and 1 μl blunting enzyme mix (NEB; # M1201AVIAL)]. Coverslips were then washed twice in 1× rCutSmart buffer (NEB; #B6004S) and twice in 1×T4 ligase buffer (NEB; # B0202S). Then, *in situ* ligation was performed overnight at 16°C in a sealed humid chamber, in 100 μl final volume per coverslip using: 2 μl T4 Ligase (NEB; #M0202S), 5 μl 10 μM biotinylated linker (5′-TACTACCTCGAGAGTTACGCTAGGGA-TAACAGGGTAATATAGTTT[biotin– dT]TTTCTATATTACCCTGTTA-TCCCTAGCGTAACTCTCGAGGTAGTA-3′), 10 μl 10× T4 Ligase Buffer (NEB; # B0202S), 1 μl dATP solution 100 mM (NEB; # N0440S), 1 μl BSA (20 mg/ml; NEB; #B9200S) and 81 μl H2O. Coverslips were washed twice in PBS and processed using a primary antibody against biotin partnered with a primary antibody directed against the protein under investigation (see Table 5).

### RNA immunoprecipitation (RIP)

Damaged and undamaged DIvA and NIH2/4 cells were lysed and RIP was performed as shown before ^6,92^. Immunoprecipitation was conducted overnight with ten μg of anti-BMI1 (Bethyl; #A301-694A) or normal rabbit IgG (Cell Signaling; #2729) as a control. DDRNAs were reverse transcribed using miScript RT II Kit (Qiagen; #1038703) and analyzed by qPCR with QuantiTect SYBR® Green PCR kit (Qiagen; #204145). DDRNA precursors were reverse transcribed and analyzed as described above.

### Cas13d RNA guides generation

Optimized guide RNAs (Table 4) against dilncRNAs were designed using the https://cas13design.nygenome.org/ software. Synthetized oligos for guide RNAs were then cloned in the crRNA cassette encoding pLentiRNAGuide_001 - hU6-RfxCas13d-DR1-BsmBI-EFS-Puro-WPRE plasmid (Addgene plasmid #138150) following the protocol by ^93^. Briefly, 5µg of the lentiviral CRISPR plasmid was digested and dephosphorylated with FastDigest BsmBI (Fermentas; # FERMFD0454) for 30 min at 37°C. The digested plasmid was purified using a QIAquick Gel Extraction Kit (Qiagen; #28506) and eluted in EB. Each pair of the synthetized oligos was phosphorylated and annealed using a 10X T4 Ligation Buffer (NEB; # B0202S), at 37°C for 30min and at 95°C for 5 min with a ramp down to 25°C at 6°C/min. The final ligation step was carried out for 10 min at room temperature using a 2X Quick Ligase Buffer (NEB; #B2200S) and a Quick Ligase (NEB; #M2200S). The generated plasmids were first validated by PCR amplification on clones with GoTaq Flexi DNA Polymerase (Promega; # M8291) and Sanger sequencing.

**Table 4:**
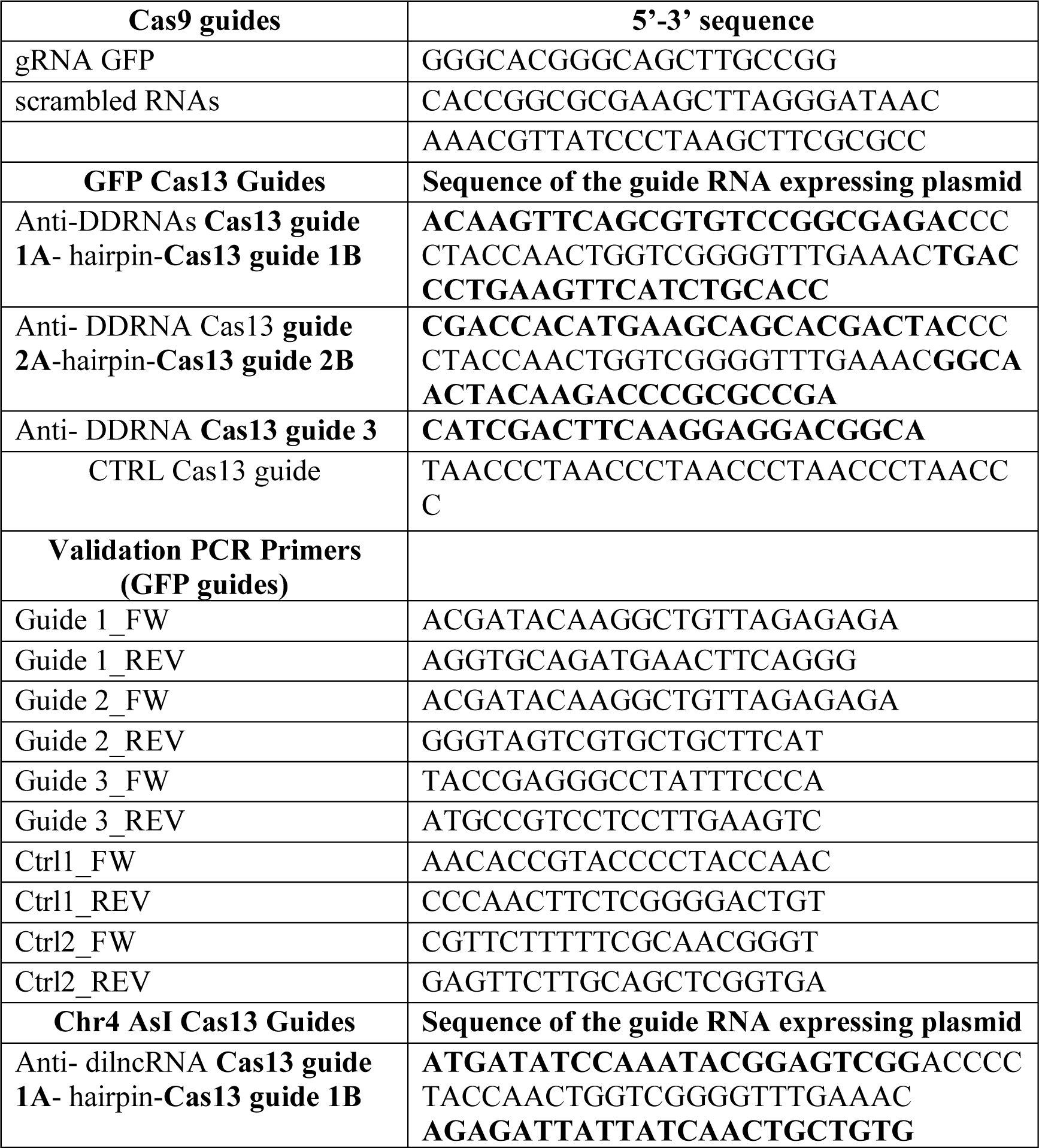
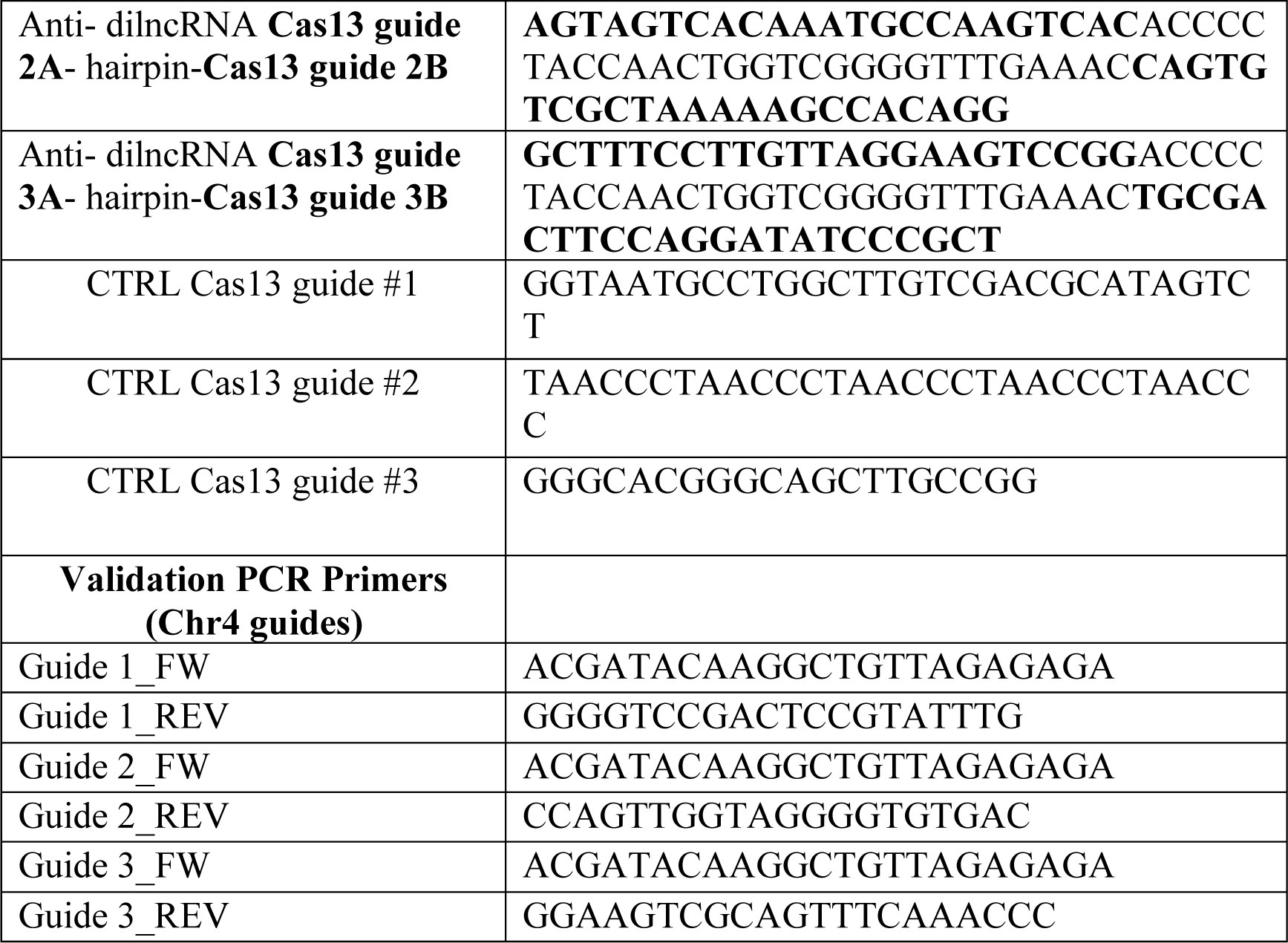
Cas13 guides and primers used in this study.

**Table 5:** Antibodies used in this study.

| **Primary Antibody** | **Host species** | **Concentration** | **Application** | **Supplier** | **Cat. N.** |
| --- | --- | --- | --- | --- | --- |
| Anti-53BP1 | goat | 1:2000 | IF | Bethyl | A303-906a |
| Anti-actin | mouse | 1:5000 | WB | Sigma-Aldrich | A5441 |
| Anti-ATM pS1981 | mouse | 1:400; 1ug/IP | IF; ChIP | Rockland | 200-301-400 |
| Anti-ATM | Mouse, clone SYR6D4 | 1:1000 | WB | Sigma-Aldrich | A6093 |
| anti-BMI1 | Rabbit, monoclonal | 1:4000 | DI-PLA | Bethyl | BLR119H |
| Anti-BMI1 | rabbit | 1:2000 | WB | Bethyl | A301-694A |
| anti-BMI1 | rabbit | 1:50 | ChIP | Cell Signaling | 6964 |
| anti-biotin | mouse | 1:4000 | DI-PLA | Sigma-Aldrich | B7653 |
| Anti- H2AX | Rabbit | 1:1000 | WB | Millipore | 07627 |
| Anti-γH2AX | mouse | 1:1000 | IF | Millipore | 05-636 |
| Anti-γH2AX | rabbit | 1:1000 | IF | Cell Signaling | 20 E3 |
| Anti-γH2AX | rabbit monoclonal | | ChIP | Abcam | Ab81299 |
| Anti-H2AK119Ub | mouse | 1:500 | IF | Millipore | 05-678 |
| anti-H2AK119Ub | mouse | 1:100 | ChIP | Millipore | 05-678 |
| Anti-β-tubulin | mouse | 1:2000 | WB | Sigma-Aldrich | T5168 |
| Anti-vinculin | mouse | 1:1000 | WB | Millipore | MAB3574 |
| anti-BMI1 | rabbit | 1:50 | ChIP | Cell Signaling | 6964 |
| anti-biotin | mouse | 1:4000 | DI-PLA | Sigma-Aldrich | B7653 |
| anti-BMI1 | Rabbit, monoclonal | 1:4000 | DI-PLA | Bethyl | BLR119H |
| Anti-Cas9 | Rabbit, monoclonal | 1: 20 000 | WB | Abcam | ab189380 |
| **Secondary Antibodies** |  |  |  |  |  |
| Anti-mouse Alexa 568 IgG | donkey | 1:400 | IF | Life Technologies | A10037 |
| Anti-rabbit Alexa 568 IgG | donkey | 1:400 | IF | Life Technologies | A10042 |
| Anti-goat Alexa 488 IgG) | donkey | 1:400 | IF | Life Technologies | A32814TR |
| anti-mouse Alexa 647 IgG | donkey | 1:400 | IF | Life Technologies | A-31571 |
| anti-mouse Alexa 647 IgM | goat | 1:400 | IF | Abcam | Ab150123 |
| anti-mouse IgG HRP | donkey | 1:10 000 | WB | Abcam | Ab97030 |
| Anti-rabbit IgG HRP | donkey | 1:10 000 | WB | Abcam | Ab97064 |

### Cell cycle distribution analysis

Following 72 hours of knock-down, DIvA cells were washed once with 1x PBS, gently trypsinised and collected in 1x PBS. Cells were then fixed in cold 100% ethanol while vortexing. Following fixation, cells were left at 4°C for at least 2 hours and were then centrifuged at 9°C for 5 min at 2000 rpm. The resulting pellet was resuspended in 1 mg/mL propidium iodide (PI) (Sigma; #P4170-100MG) + 10 mg/mL RNAse A (Sigma-Aldrich; #10109142001) in 1x PBS and incubated at 37° C for 30 min. Samples were analyzed on an S3e Cell sorter (BioRad) and results were plotted using the FlowJo 10.8 software. 10^4^ events were analyzed for each condition.

### Statistical analysis

Prism 9 software (GraphPad) was used to generate graphs, to perform statistical analysis and to remove outliers with the Robust regression and Outlier removal method in figures SI, Figure 3H and figures S3D,F,H. Statistical analysis was performed with the One-way ANOVA test, unless differently indicated.

## Acknowledgments

We thank Prof. R.A. Greenberg (Department of Cancer Biology, Abramson Family Cancer Research Institute, Philadelphia, USA), Dr. Andrea Ventura (Cancer Biology and Genetics Program, Memorial Sloan Kettering Cancer Center, New York, United States), Dr. E. Soutoglou (Genome Damage and Stability Centre, Sussex University, School of Life Sciences, University of Sussex, Brighton, BN1 9RH, UK), Dr. Guido Drexler (Ludwig-Maximilians-University of Munich, LMU, Department of Radiation Oncology, Munich, Germany), Dr. Martin Mistrik (Palacký University Olomouc, Institute of Molecular and Translational Medicine, Olomouc, Czech Republic) and Dr. Gaëlle Legube (CBI, MCD, Univsersité Paul Sabatier, Toulouse, France) for providing crucial reagents. We thank Dr. Alexandra Mancheno for help with bioinformatic competences and RNA sequencing data deposition in GEO. We also thank Dr. Simone Sabbioneda and Dr. Anna Garbelli for the help in cell cycle experiments. The research was funded by EPIGEN, Progetto Bandiera Epigenomica; Fondazione Cariplo project 2014-1215; The Association for International Cancer Research (AICR)-World Wide Cancer Research (WWCR)-rif. N.14-1331. S.F. is supported by AriSLA (projects “DDRNA&ALS” 2018-2020 and its follow up “DDR&ALS” 2021-2024) and by a grant POR FESR 2014-2020 Regione Lombardia (InterSLA project) PNRR-National Center for the development of gene therapy and drugs with RNA technology CN3; Parternariato esteso PNRR “AgeIT”. S.M. is supported by IUSS Pavia. F.E. is supported by an AIRC fellowship for Italy. F.d’A.d.F laboratory is supported by: Progetti di Ricerca di Interesse Nazionale (PRIN) 2015 “ATR and ATM-mediated control of chromo-some integrity and cell plasticity”; Progetti di Ricerca di Interesse Nazionale (PRIN) 2017-2017NWEXEP “RNA and genome Instability”; Progetti di Ricerca di Interesse Nazionale (PRIN) 2020-2020CXFL4T “Interplay between genotoxic and mechanical stress in neurodegeneration”; PRIN 2022 2022R7LH5T “The impact of telomere dysfunction in healthspan and lifespan in wild-type mice; ERC advanced grant (TELORNAGING—835103); AIRC-IG (21762); Telethon (GGP17111); AIRC 5X1000 (21091); ERC PoC grant (FIREQUENCER—875139); FRRB—Fondazione Regionale per la Ricerca Biomedica—under the frame of EJP RD, the European Joint Programme on Rare Diseases with funding from the European Union’s Horizon 2020 research and innovation program under the EJP RD COFUND-EJP NO 825575.

## Author contribution

SF conceived the study and designed the experiments. FdAdF funded part of the study and contributed in defining the publication strategy. FI performed the bioinformatic analyses of RNA sequencing in Figure 1A and edited the relative part in the result’s section. AC performed bioinformatic analysis during the revision process. AR performed part of the biological replicates for panels in Figure S4J and S5I and J. UG performed RIP analyses for Bmi1 in NIH2/4 and DiVA cellular systems (Figures 5G,H and S4H-K), wrote and edited the manuscript. A.dL. generated the HeLa GFP cellular system and S.T. designed and preliminary tested ASOs toxicity in the Hela GFP system. M.C. generated the inducible Cas13d HeLa-GFP cellular system. L.M. prepared the samples for Figure 2A and performed the experiments in Figures 2B, 4H-I, 5D, and S4C-D. SM performed the experiments in figures 2C, 3F-H, S2H, S3, S5E, assembled figure legends and material and methods and edited the text. F.E. prepared the samples for Figure 2A and performed the experiments in figures 1B-E, 2B, 2D-E, 3B-G, 4G, 5E-F, S2D-G and S5H-I, assembled figure legends and material and methods, wrote and edited the text. IC perform all the remaining experiments. SF and FE wrote the text of the manuscript and the multiple round of responses to referee; FdAdF, UG, SM edited the texts.

## Notes

### Competing Interest Statement

The authors have declared no competing interest.

