## Supplementary Figures for "DROSHA, DICER and Damage-Induced long ncRNA control BMI1-dependent transcriptional repression at DNA double-strand break"

**Figure supplementary 1.**

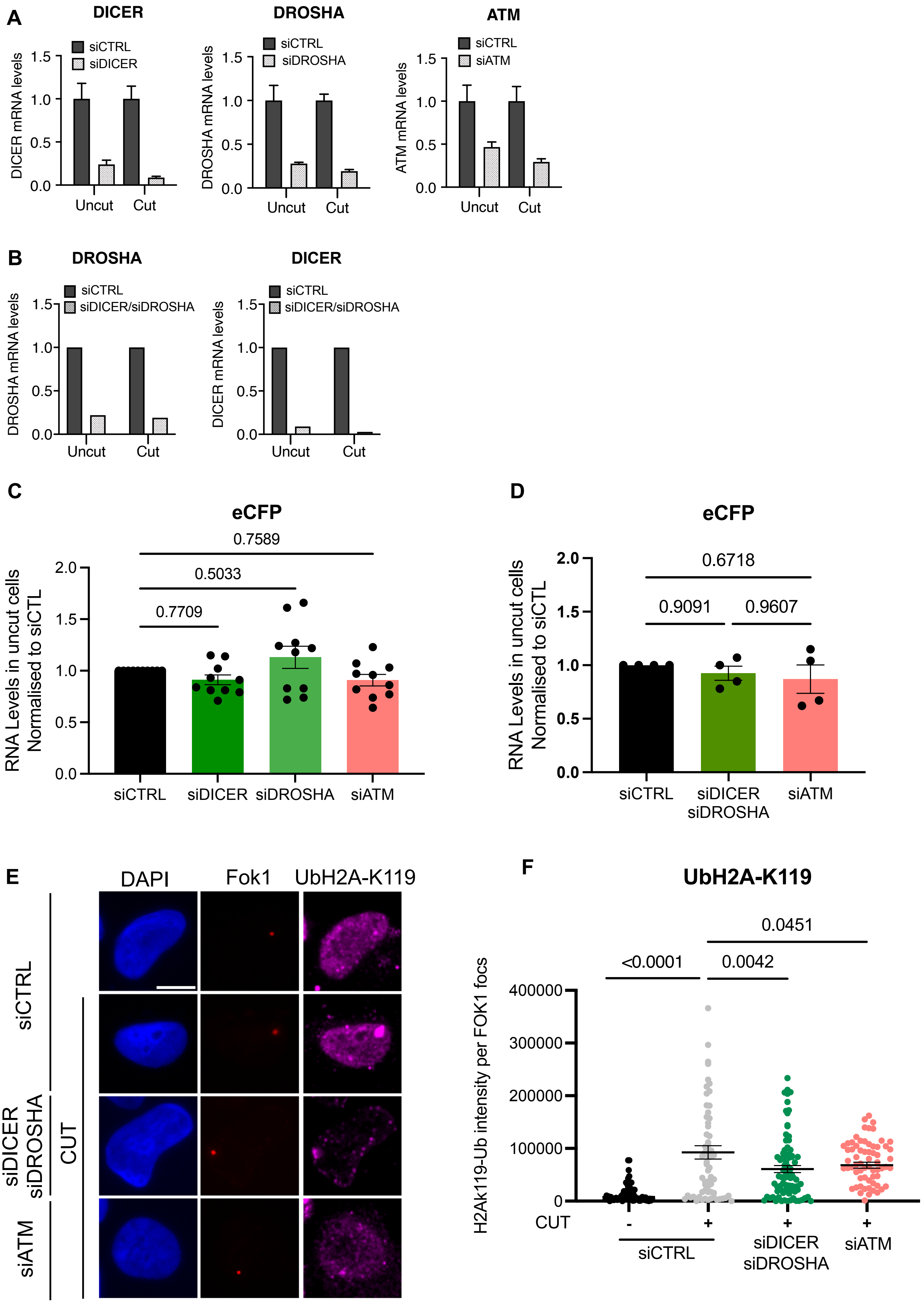

**Figure Supplementary 1. A.** Representative RT-qPCR analysis of DICER, DROSHA and ATM mRNA levels in both cut and uncut U2OS 2-6-3 cells treated with non-targeting control siRNAs (siCTRL) or siRNAs against DICER, DROSHA or ATM transcripts (siDICER, siDROSHA and siATM, respectively). **B.** Representative RT-qPCR analysis of DROSHA and DICER mRNA levels in both cut and uncut U2OS 2-6-3 cells treated with siDICER and siDROSHA. **C.** RT- qPCR analysis of CFP mRNA levels in both cut and uncut U2OS 2-6-3 cells treated with siDICER, siDROSHA and siATM. Statistical analyses in panels were performed by One-Way ANOVA. ns: not significant. **D.** RT- qPCR analysis of CFP mRNA levels in both cut and uncut U2OS 2-6-3 cells treated with siDICER and siDROSHA. **E.** Images of ubH2A-K119 signal at the Cherry-Lac locus in uncut and cut U2OS 2-6-3 cells transfected with siCTRL, siDICER and siDROSHA, or siATM. **F.** Quantification of the intensity of ubH2A-K119 signal forming at FOK1 focus in cells treated as in C. Error bars represent SEM of more than 50 cells from three independent experiments. Statistical analyses in panels C, D and F were performed by One-Way ANOVA.

### Figure Supplementary 2.

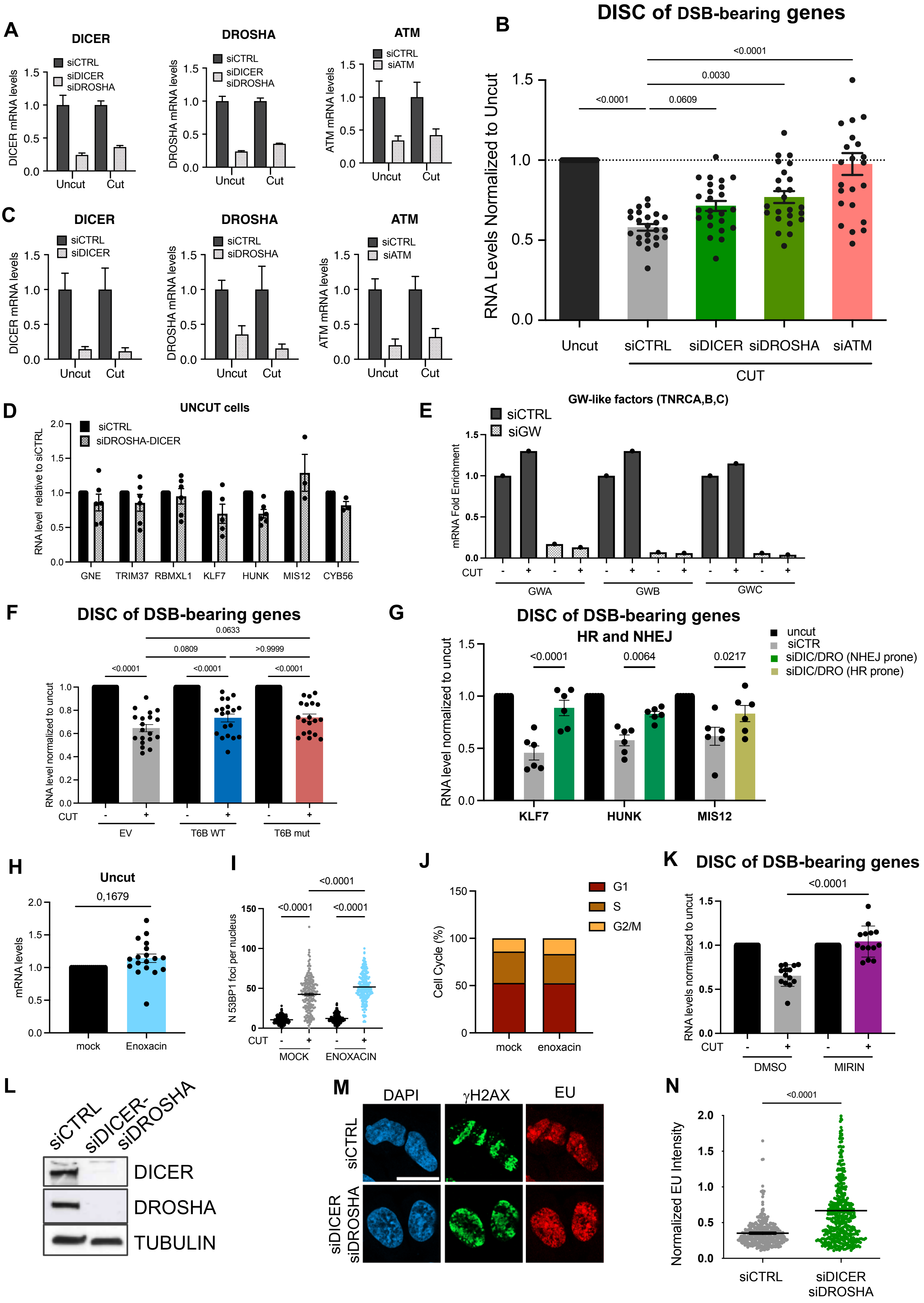

**Figure supplementary 2: A.** Representative RT-qPCR analysis of DICER, DROSHA or ATM mRNA levels in both cut and uncut DIvA cells treated with non-targeting control siRNAs (siCTRL) or siRNAs against DICER, DROSHA or ATM transcripts (siDICER, siDROSHA and siATM, respectively). **B.** RT-qPCR analysis of break-bearing gene expression in cut versus uncut cells treated with siCTRL, siDICER, siDROSHA or siATM. Data are relative to uncut cells for each knockdown condition. Error bars represent SEM of the expression of the selected 6 genes from three independent experiments. **C.** Representative RT-qPCR analysis of DICER, DROSHA or ATM mRNA levels in both cut and uncut DIvA cells treated with non-targeting control siRNAs or siRNAs against DICER, DROSHA or ATM transcripts. **D.** The expression of the selected genes was analysed by RT-qPCR in uncut cells transfected with siCTRL, or siDICER and siDROSHA, and shown individually. Data are relative to the uncut siCTRL condition. **E.** Representative RT-qPCR analysis of TNRCA, TNRCB or TNRCC mRNA levels in both cut and uncut DIvA cells, treated with siCTRL or siRNAs against TNRCA, TNRCB or TNRCC transcripts. **F.** RT-qPCR analysis of break-bearing gene expression in cut versus uncut cells transfected with the empty vector (EV), wild type T6B (T6B WT) or mutant T6B peptide (T6B mut). Data are relative to uncut cells for each transfection condition. Error bars represent SEM of the expression of the 6 genes from three independent experiments. **G.** RT-qPCR analysis of the expression of NHEJ- (KLF7, HUNK) and HR-prone (MIS12) genes in cut versus uncut cells, treated with siCTRL, siDICER, siDROSHA or siATM. Data are relative to uncut cells for each knockdown condition. **H.** RT-qPCR analysis of break-bearing genes expression in uncut cells treated or not with enoxacin. Data are relative to untreated uncut cells. Error bars represent SEM of the expression of the selected 6 genes from three independent experiments **I.** Quantification of 53BP1 foci in cut and uncut cells treated or not with enoxacin. Error bars represent SEM of more than 150 cells from two independent experiments. **J.** Cell cycle analysis of DiVA cells treated or not with enoxacin and stained with propidium iodide. Percentage of cells in cell cycle phases is indicated for each condition. Ten thousand cells were analyzed for each condition. **K.** RT-qPCR analysis of the mRNA levels of the 6 genes relative to uncut, in DMSO or MIRIN treated cells. Data are relative to uncut cells for each treatment condition. Error bars represent SEM of the expression of the six genes from two independent experiments. **L.** Representative immunoblot of DROSHA and DICER protein levels in DIvA cells transfected with siCTRL, siDICER and siDROSHA. Tubulin was used as a loading control. **M.** Images of γH2AX stripes and EU incorporation in laser micro-irradiated DIvA cells, transfected with siCTRL or siDICER and siDROSHA. DNA was counterstained with DAPI. Scale bar: 10 μm. **N.** Quantification of EU intensity normalized on γH2AX levels in laser micro-irradiated U2OS cells treated as in M. Error bars represent SEM of more than 200 cells from three independent experiments. Statistical test used was Student t-test. Statistical analyses in panels B, F, G and H were performed by One-Way ANOVA.

### Figure supplementary 3.

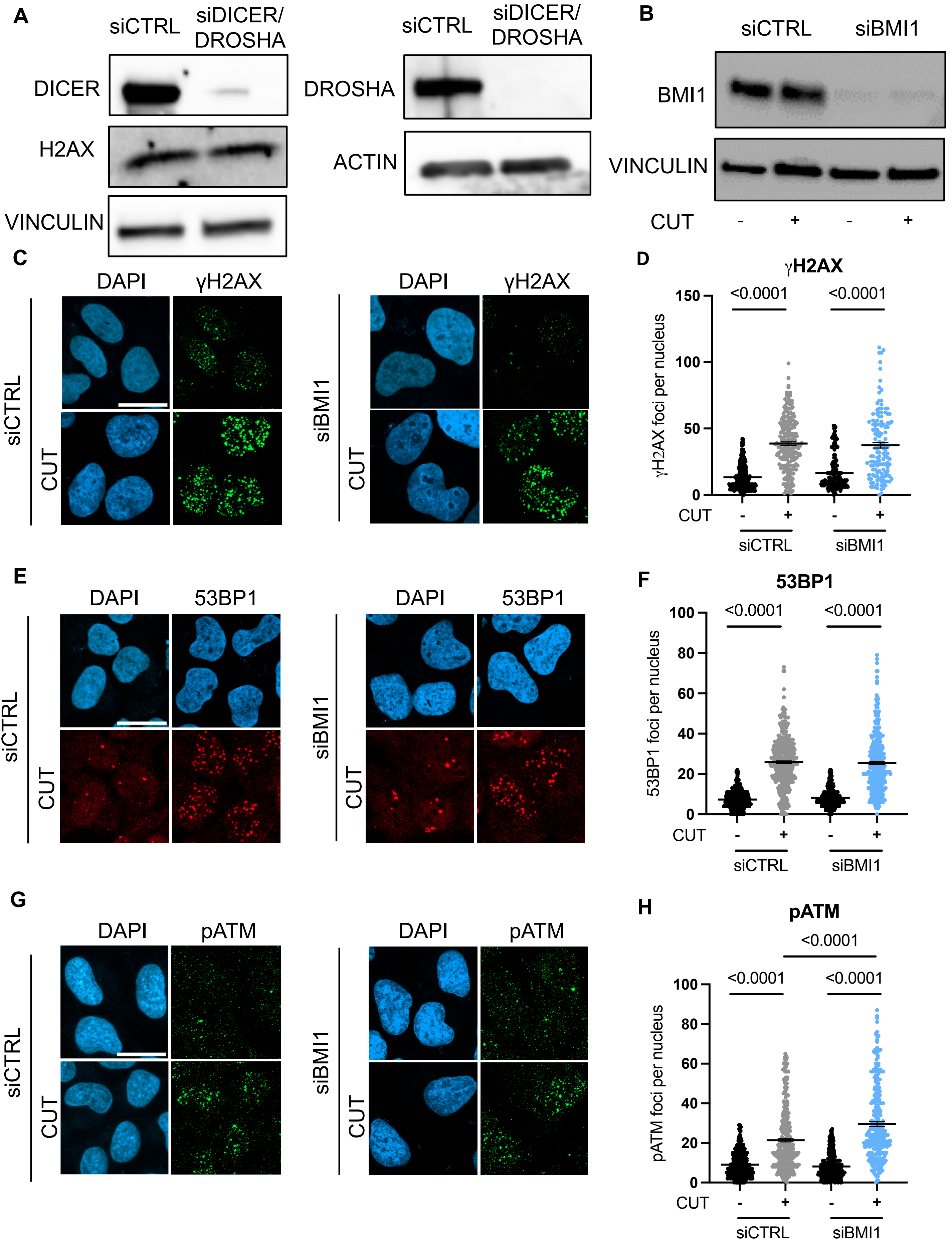

**Figure supplementary 3: A, B.** Immunoblot analysis of DICER, DROSHA, BMI1 and H2AX protein levels in uncut DIvA cells transfected with non-targeting siRNAs (siCTRL), siRNAs against DICER and DROSHA (siDICER/DROSHA) or BMI1 transcript (siBMI1). Vinculin or Actin were used as loading controls. **C.** Images of cut and uncut DIvA cells transfected with siCTRL or siBMI1 and stained for γH2AX. **D.** Quantification of γH2AX foci as determined in C. **E.** Images of 53BP1 foci from cells treated as in C. **F.** Quantification of 53BP1 foci as determined in E. **G.** Images of cells treated as in C and stained for pATM. DNA was counterstained with DAPI in C, E and G. Scale bars in C, E and G: 20 μm. **H.** Quantification of pATM foci as determined in G. Error bars in D, F and H represent SEM of more than 150 cells from three independent experiments. Statistical analyses in panels D, F and H were performed by One-Way ANOVA.

### Figure supplementary 4.

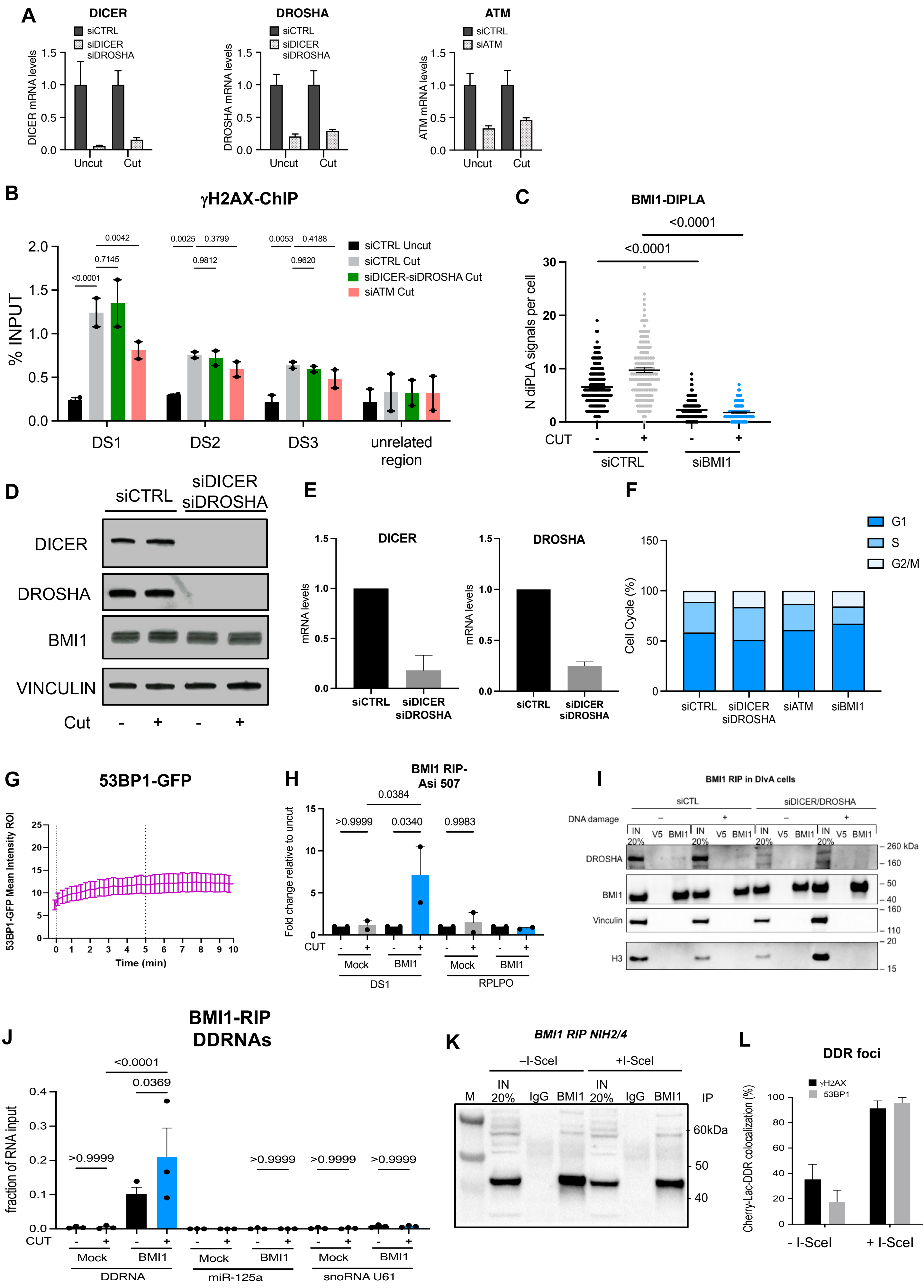

**Figure Supplementary 4. A.** Representative RT-qPCR analysis of DICER, DROSHA or ATM mRNA levels in both cut and uncut DIvA cells treated with non-targeting siRNAs (siCTRL), siRNAs against DICER, DROSHA or ATM transcripts (siDICER, siDROSHA and siATM, respectively). **B.** ChIP-qPCR analysis for γH2AX at three AsiSI sites and an unrelated region, performed in cut and uncut cells treated with siCTRL, siDICER/siDROSHA, or siATM. Locations of AsiSI break sites are as in Figure 4A. Error bars represent SEM from three independent experiments. Statistical analyses were performed by One-Way ANOVA. **C.** Quantification of DI-PLA signals for BMI1 recruitment to sites of DNA damage in cut and uncut cells treated with siCTRL or siRNAs against BMI1 transcript (siBMI1). Error bars represent SEM of more than 200 cells from three independent experiments. Statistical analyses were performed by One-Way ANOVA. **D.** Immunoblots of DICER, DROSHA and BMI1 protein levels in cut and uncut DIvA cells transfected with siCTRL or siDICER and siDROSHA. Vinculin was used as a loading control. **E.** Representative RT-qPCR analysis of DROSHA and DICER mRNA levels in U2OS cells transfected with siCTRL or siDICER and siDROSHA. **F.** Cell cycle analysis of DiVA cells treated with siCTRL, siDICER and siDROSHA, siATM or siBMI and stained with propidium iodide. Percentage of cells in cell cycle phases is indicated for each condition. More than ten thousand cells were analyzed for each condition. **G.** Quantification of 53BP1-GFP signal intensity at laser tracks over time in U2OS cells. Values are relative to GFP intensity prior to irradiation; error bars represent SEM of at least 12 cells from three independent experiments. Statistical analyses in panels B and C were performed by One-Way ANOVA. **H.** RNA immunoprecipitation (RIP) of BMI1-associated RNAs followed by RT-qPCR analysis of dilncRNAs generated at DS1, performed in uncut and cut DIvA cells. RPLP0 was used as a negative control. **I.** Immunoblot of DROSHA, BMI1, Vinculin and histone H3 from protein extracts of RIP experiments described in Figure 4H and performed in uncut and cut DIvA cells. **J.** RNA immunoprecipitation (RIP) of BMI1-associated RNAs followed by RT-qPCR analysis of DDRNAs produced at the exogenous LAC-operon locus in uncut and I-SceI-cut NIH2/4 cells; RPLP0, miR-125a and snoRNA U61 were used as negative controls. Error bars represent SEM from three independent experiments. Statistical analysis was performed using Two-Way ANOVA. **K**. Immunoblot of BMI1 from protein extracts of RIP experiments performed in uncut and I-SceI-cut NIH2/4 cells. L Percentage of colocalization of γH2AX (black bars) and 53BP1 (grey bars) in uncut and I-SceI-cut NIH2/4 cells relative to Figure 4H.

### Figure Supplementary 5.

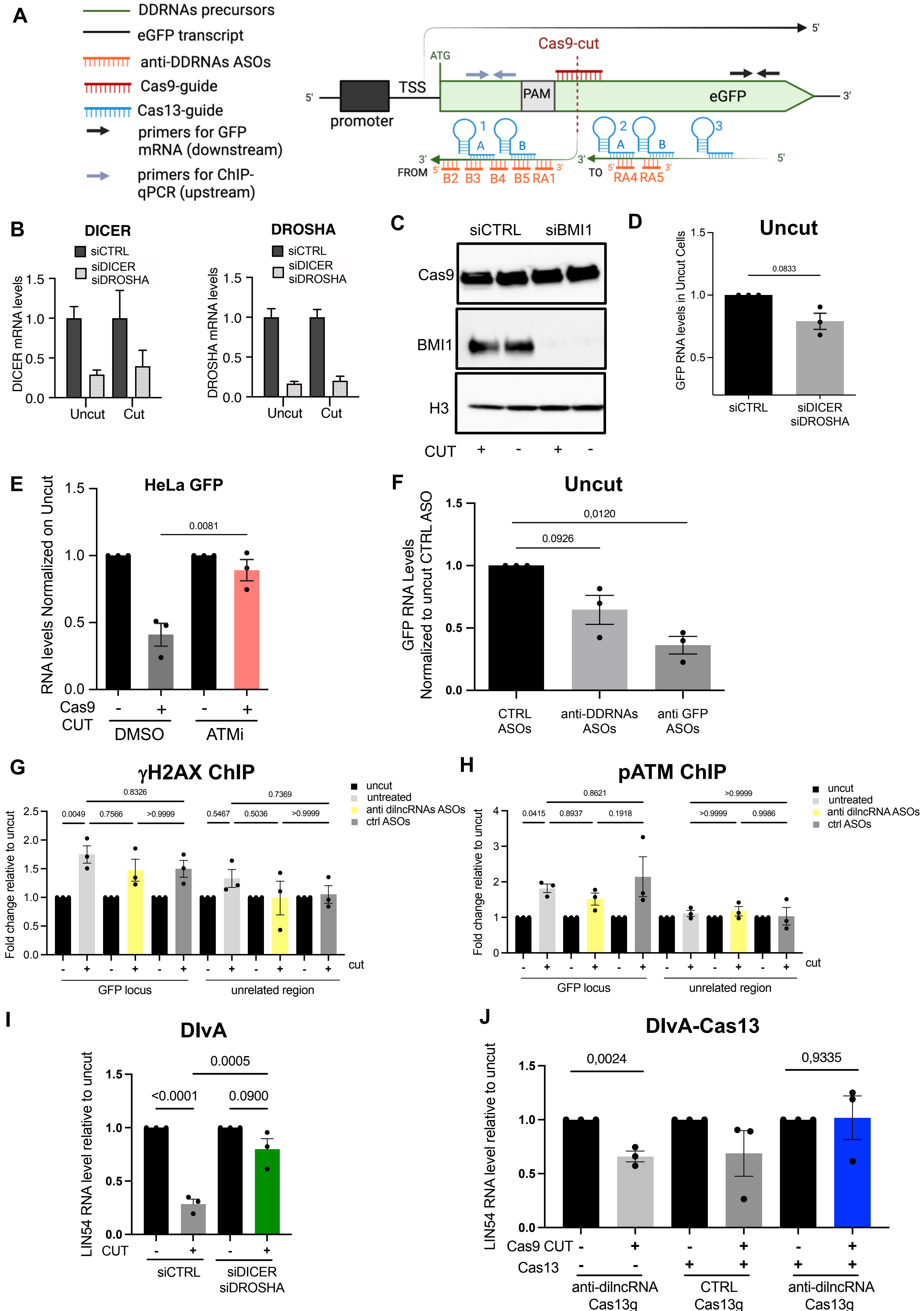

**Figure Supplementary 5: A.** Schematic representation of the GFP locus in the HeLa-GFP cellular system. The eGFP gene (long black arrow) is transcribed from the transcription start site (TSS), localized downstream the promoter. From the guide-directed Cas9 cut, sense and antisense damage-induced long non-coding RNAs (dilncRNAs, DDRNAs precursors) are generated both from and towards the cut site. In figure, only antisense dilncRNAs are represented (green arrow). Seven different ASOs sequences (orange) were used to target antisense dilncRNAs: five upstream the cut site (B2-B5 and RA1), and two downstream the cut site (RA4 and RA5). ASO sequences do not overlap with the Cas9-guide sequence or with each other. All ASOs were co-transfected to achieve DDRNAs targeting. Similarly, two Cas13 guide sequences (guide 1A and 1B), interspaced by the short hairpin necessary for guide RNA recognition, were designed upstream and three (guides 2A, 2B and guide 3) were designed downstream the break. Plasmid containing guide 1A and 1B, 2A and 2B and 3 were co-transfected to target DDRNAs precursors. A pair of primers for ChIP-qPCR the analysis was designed upstream the cut (short grey arrows), while a pair of primers to evaluate the eGFP transcript level was designed downstream the cut (short black arrows). For a complete list of ASOs, RNA guides and primers see Tables 1, 3 and 4 in the methods section. The figure was created with BioRender.com. **B.** Representative RT-qPCR analysis of DROSHA and DICER mRNA levels in both cut and uncut HeLa cells treated with non-targeting control siRNAs (siCTRL) or siRNAs against DROSHA and DICER transcripts (siDICER/siDROSHA). **C.** Immunoblot of Cas9 and BMI1 protein levels in scramble-guide-Cas9 (uncut) and GFP-guide-Cas9 (cut) transfected HeLa-GFP cells, treated with siCTRL or siBMI1. Histone H3 was used as a loading control. **D.** RT-qPCR analysis of GFP mRNA levels in uncut HeLa-GFP cells transfected with siCTRL or siDICER and siDROSHA. Error bars represent SEM from three independent experiments. Statistical test used was paired Student t-test: ns: not significant. **E.** RT-qPCR analysis of GFP mRNA levels in cut and uncut HeLa-GFP cells treated with DMSO or ATM inhibitor ATMi). Error bars represent SEM from three independent experiments. **F.** RT-qPCR analysis of GFP mRNA levels in uncut cells treated with control, DDRNA-targeting ASOs (GFP antisense transcript) or GFP sense transcript-targeting ASOs. Error bars represent SEM from three independent experiments. **G, H.** ChIP-qPCR analysis for γH2AX (G) and pATM (H) at the GFP locus and at an unrelated region, performed in uncut (scramble-guide-Cas9) and cut (GFP-guide-Cas9) cells treated with control or dilncRNAs-targeting ASOs. Data are relative to uncut cells after the subtraction of mock values for each treated condition. Error bars represent SEM from three independent experiments. **I.** RT-qPCR analysis of LIN54 (DSB at Chr4) expression in cut and uncut DIvA cells treated with siCTRL, siDICER and siDROSHA. Data are relative to uncut cells for each knockdown condition. Error bars represent SEM from three independent experiments. **J.** RT-qPCR analysis of LIN54 (DSB at Chr4) mRNA levels in cut and uncut DIvA cells stably expressing inducible Cas13. Cells were treated with a mix of the three control Cas13 RNA guides (CTRL Cas13g) or guides targeting dilncRNAs generated downstream the AsI-SI cut site on Chr4 (Chr4 RNA Cas13g). Error bars represent SEM from three independent experiments. Statistical analysis in panel D was performed by unpaired t test. Statistical analyses in panels E, F, G, H, I and J were performed by One-Way ANOVA.
